# Water migration through enzyme tunnels is sensitive to choice of explicit water model

**DOI:** 10.1101/2023.08.14.553223

**Authors:** Aravind Selvaram Thirunavukarasu, Katarzyna Szleper, Gamze Tanriver, Karolina Mitusinska, Artur Gora, Jan Brezovsky

## Abstract

Understanding the utilization of tunnels and water transport within enzymes is crucial for the catalytic function of enzymes, as water molecules can stabilize bound substrates and help with unbinding processes of products and inhibitors. Since the choice of water models for molecular dynamics simulations was shown to determine the accuracy of various calculated properties of the bulk solvent and solvated proteins, we have investigated if and to what extent the water transport through the enzyme tunnels depends on the selection of the water model. Here, we have focused on simulating enzymes with various well-defined tunnel geometries. In a systematic investigation using haloalkane dehalogenase as a model system, we focused on the well-established TIP3P, OPC, and TIP4P-Ew water models to explore their impact on using tunnels for water molecules transport. The TIP3P water model showed significantly faster migration, resulting in the transport of approximately 2.5 times more water molecules in comparison to OPC and 2.0 times greater than the TIP4P-Ew. The increase in migration of TIP3P water molecules was mainly due to faster transit times, and in the case of narrower tunnels, greater concurrent transport was evident as well. We have observed similar behavior in two different enzymes with buried active sites and different tunnel network topologies, indicating that our findings are likely not restricted to a particular enzyme family. Our study emphasizes the critical importance of water models in comprehending the use of enzyme tunnels for small molecule transport. Given the significant role of water availability in various stages of the catalytic cycle and solvation of substrates, products, and drugs, choosing an appropriate water model might be crucial for accurate simulations of complex enzymatic reactions, rational enzyme design, and predicting drug residence times.

## Introduction

Water molecules play crucial roles in the structure, function, and stability of proteins and other biomolecules,^1,2^ They often form intricate hydrogen bond networks and thereby stabilize the hydrophilic cavities, protein subunits etc., for example, the hexametric hydrophilic cavity of insulin is maintained by approximately ten water molecules which form intricate hydrogen bond networks.^3^ Water aids in protein folding and markedly influences the dynamics of the hydrated molecules.^4,5^ Waters not only mediate the interaction of residues in the short-range but also influence the allosteric regulation of proteins.^6,7^ They also play a role in ligand binding by creating a favorable environment for forming H-bonds, stabilizing the internal cavities of the protein.^8–11^. In proteins where such internal cavities are buried deep in their structure, the tunnels connecting them with their exterior are required to transport solvents, ions, and cognate molecules, e.g., substrates and products of enzymatic catalysis.^12–16^ The selectivity of these functional tunnels is controlled by molecular mechanisms like molecular gates which play an important role in substrate and product release. ^17–21^

The unquestionable role of the water molecules in biological systems is reflected by number of various tools and methods both including water in systems description and utilizing them as an important tool.^22^ As an example, explicit water molecules become an essential component of the majority of atomic-level molecular dynamics (MD) simulations.^10,11^ The behavior of proteins in MD simulations is dependent on the models of waters and the force fields being used in the simulations.^23–25^ In intrinsically disordered proteins (IDPs), water plays a crucial role by interacting with the protein backbone and sidechains, mediating interactions between different protein regions.^26–28^ Comparison of properties of five IDPs simulated with different water models revealed significant discrepancies among the simulations as well as disagreement with experimental data.^29^ To partially address this discrepancy, a water model needed to be adjusted^29^ and the force field to simulate folded and unfolded states developed.^30^ Similarly, a recent study observed differences in the duration of open states of tunnels in three variants of haloalkane dehalogenase LinB between simulations using OPC and TIP3P models.^31^ Finally, the trafficking of water molecules by aquaporin AQP1, which can perform such tasks with large diffusivity,^32^ was shown to be dependent on the water models employed.^33^ In this work, the simulations with three water models (TIP3P, TIP4P/2005, and OPC) revealed significant differences in their osmotic permeabilities via AQP1.^33^

Here, we delve into the impact of three prominent water models on the transportation of water through enzyme tunnels. Such tunnels are structurally and functionally distinct from aquaporin channels since they need to compromise for the trafficking of substrate/product as well as water molecules. We have chosen haloalkane dehalogenase DhaA (DhaA) from *Rhodococcus rhodochrous*,^34^ as our model system to examine the influence of water models on water migration under a range of well-controlled tunnel geometries, benefiting from the availability of recent high-throughput simulations data on tunnel network in this enzyme (**Figure 1A**). ^35^ We then verified our findings with two other distinct enzyme systems, i.e., alditol oxidase (AldO) from *Streptomyces coelicolor A3*,^36^ and cytochrome P450 2D6(Cyt P450) (**Figure 1A**).^37^ From the variety of available water models, we have chosen the following three: TIP3P, TIP4P, and OPC (**Figure 1B**). TIP3P is still widely employed in simulations often due to its computationally low cost and well-known behavior in a variety of systems. OPC and TIP4P, on the other hand, were chosen due to their ability to more accurately reproduce the characteristics and behavior of bulk water (**Figure 1C**).^38,39^ The employed protocol allowed us to probe the details of mechanisms behind the water transport through these tunnels, including the number of transported water molecules, required transit times, parallelism of transport, and the likelihood of tunnels adopting conductive states enabling water migration.

**Figure 1:**
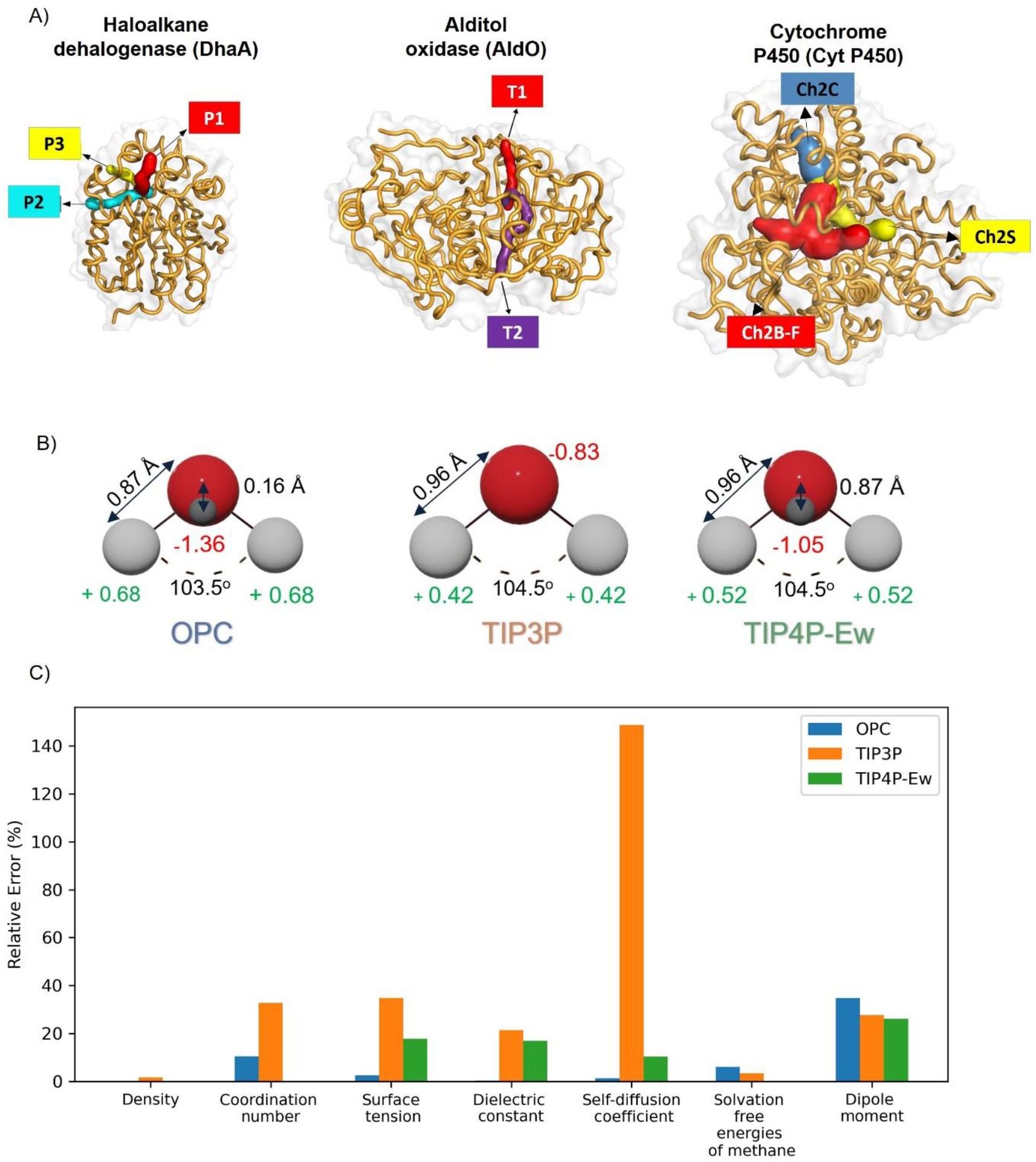
Overview of studied systems and water models. (A) Structures of three selected enzymes with their main tunnels are shown. (B) The arrangement of water molecules for the three studied water models. (C) Errors in the key physiochemical properties of the studied water models relative to the experimental data.

## Materials and Methods

### Extraction of haloalkane dehalogenase DhaA conformations with defined tunnel geometries

The starting structures were systematically selected based on their bottleneck radii (radius of the narrowest part of the tunnel) of the main P1 tunnel, which has been shown to constitute the dominant water conduit in previously conducted high-throughput simulations of this enzyme.^35^ We have also included tunnels with bottleneck radii starting from 1 Å to reflect upon the recent observation of significant transport of water molecules even through the tunnels with bottleneck radii lower than 1.4 Å.^35^ Overall, in our investigation, we defined five distinct Tunnel Conformational Groups (TCGs) with increasing bottlneck radii of P1 tunnel. Here, each TCG consists of a collection of tunnels sampled in five independent simulations restrained at conformations with equivalent bottleneck radii of these tunnels. Such systematic grouping enables exploration of the dynamic water behavior occurring with these well-defined tunnel geometries. The details of the bottleneck radii of the selected starting frames are available in **Table S1**.

### Preparation of selected frames for comparative MD simulations with different water models

The selected frames were extracted from the trajectories as PDB files to maintain the positions of the pre-equilibrated water molecules from the previous simulations. Using *tLeap* module of AmberTools21,^40^ the OPC, TIP3P, and TIP4P-Ew models for water molecules and ff19SB for proteins were set, and AMBER-formatted topology and input coordinates were saved. The original dimension of the truncated octahedral box from previous simulations was enforced by *ChBox* and *cpptraj*^41^ modules of AmberTools21 for coordinates and topology, respectively. To enable the time step to 4 fs, the hydrogen mass repartitioning^42^ was performed by *parmed*^43^ module of AmberTools21.^33^

### Restrained MD simulations to generate tunnel ensembles with defined geometries

Using PMEMD and PMEDD.CUDA modules of Amber 20,^40,44^ the systems were first energy minimized by 100 steepest descent steps followed by 400 conjugate gradient steps in five rounds with decreasing harmonic restraints as follows; 500 kcal mol^−1^ Å^−2^ applied on all heavy atoms of protein, followed by 500, 125, 25, 0.0001 kcal mol^−1^ Å^−2^ applied only to the backbone heavy atoms. After the minimization, the systems were equilibrated in four stages. First, the system was gradually heated from 0 to 200 K in 20 ps NVT simulations while harmonic restraints of 5 mol^−1^ Å^−2^ were applied on the heavy atoms of protein, followed by 1 ns NVT simulations to reach the target temperature of 310 K, still maintaining the harmonic restraint of 5.0 kcal mol^−1^ Å^−2^ on the heavy atoms of the protein. The temperature was controlled with a Langevin thermostat using a collision frequency of 2.0 ps^-1^. In the third stage, 1 ns NPT simulations were performed at 310 K using a Langevin thermostat and maintaining a constant pressure of 1.0 bar using a Monte Carlo barostat while keeping 1.0 kcal mol^−1^ Å^−2^ restraints on the protein backbone atoms. The last equilibration stage consisted of an NPT run of 1 ns with a weaker restrain of 0.5 kcal mol^−1^ Å^−2^ applied on the backbone atoms of the protein. Finally, 300 ns of NPT simulations were run at 310 K maintained using a weak-coupling Berendsen thermostat and a Monte Carlo barostat with weak restraint of 0.5 kcal mol^−1^ Å^−2^ applied on the backbone atoms of the protein to maintain the tunnel geometries. The trajectories were saved at 10 ps intervals. All the simulations employed periodic boundary conditions with particle mesh Ewald method,^45^ and SHAKE^46^ algorithm, to constrain the movement of bonds involving hydrogen atoms.

### Analysis of tunnel network and migration of water molecules through the network

For all the trajectories, the geometric tunnels were calculated using CAVER 3.0^47^ with the starting points for the calculation defined by the following atoms: D106-CG, W107-CD2, F168-N, and L246-N, using a probe radius of 0.9 Å, shell radius 3 Å and shell depth 4 Å. The obtained tunnels were clustered by hierarchical clustering with a clustering threshold of 3.5 and limiting the maximum output clusters to 50 to discard less relevant tunnel clusters. Then, the movement of water from the bulk solvent to the core of the protein to the active site was traced using AQUA-DUCT version 1.0.11^48^ with the scope defined as protein and object defined as the sphere of 6 Å formed by the same atoms used for the starting point of tunnel calculation, and the other settings left at their default values. Both methods gives different and complementary analysis of tunnels network,^49^ and due to incompatible definition of the tunnels, the traced water pathways were assigned to the tunnel clusters using TransportTools version 0.95^50^. TransportTools superimposed the results of CAVER and AQUADUCT outputs from individual simulations then assigning the traced water molecules to the calculated tunnels providing a unified view of water transport events per tunnel clusters. The clustering of tunnels across the simulations was done using hierarchical clustering with complete linkage and 1.5 clustering cutoff. For the cases where the same water transport event is assigned to multiple tunnel clusters, it was resolved by using exact matching, where the tunnel geometry in the original trajectory will be used to assign the waters to respective tunnel clusters. Using a comparative analysis of TransportTools, the results of water transport statistics for user-defined groups of simulations with different water models and tunnel geometries were obtained. The exact matching analysis was performed to analyze and visualize frames where the waters are present inside the tunnel. The other settings were left default.

### Water transport in tunnels of different enzymes – cases of AldO and Cyt P450

To extend our analyses to other enzyme systems, we have performed MD simulations of AldO and Cyt P450 to generate ensembles of tunnels with narrow (TCGnarrow) and wide (TCGwide) bottlenecks geometries.

### Preparation of starting structures and their initial MD simulation

The crystallographic structures for AldO (PDB ID: 2vft) and Cyt P450 (PDB ID: 3tbg) were obtained from the Protein Data Bank.^51^ The structures were processed by removing all non-protein molecules except crystallographic waters and cofactors (FAD in AldO and HEM in Cyt P450). The protonation states of both structures were determined using H++ 3.0 webserver,^52^ at pH 8.5. For both systems, the initial waters were solvated using a tandem approach based on 3D Reference Interaction Site Model Theory (3D-RISM). ^53^ Then, using the tLeap module of Amber20^54^, each protein was placed at the center of a periodic truncated octahedral box 10 Å away from the edges. The system was solvated with an OPC water model^38^. Na^+^ and Cl^-^ ions were added to neutralize the system and reach the salt concentration of 0.1% of water. For the cofactors, General Amber Force Field (GAFF2), ^55^ was used to generate force field parameters and AM1/BCC charges utilizing the ACEPYPE script and Antechamber module of Amber20.^56^ For HEME cofactor, MCPB.py module of Amber20 was used for its parametrization.^57^ To enable the time step to 4 fs, the hydrogen mass repartitioning ^42^ was performed by *parmed*^43^ module. The prepared structures were energy minimized in five rounds with decreasing harmonic restraints and equilibrated in four stages as described for DhaA above. Finally, unrestrained 250 ns of NPT simulations were run at 310 K maintained using a weak-coupling Berendsen thermostat and a Monte Carlo barostat while saving the trajectories in 2 ps intervals.

### Extraction of frames for the defined tunnel geometries

The last 50 ns of the resulting trajectories were analyzed by CAVER to identify the tunnel networks in these enzymes, using the following atoms to define starting points for the calculation: S106-CA, E320-CA, R322-CA, H343-CA, T345-CA, and K375-CA for AldO, and L110-CA, F112-CA, L216-CA, E218-CA, Q244-CA, F247-CA, and D301-CA for Cyt P450. For AldO, the frames were selected according to the bottleneck radii of the main tunnel (T1) of 1.0 Å and 2.0 Å to produce TCGnarrow and TCGwide (**Table S2**). For Cyt P450, the frames were selected according to the bottleneck radii of 1.0 Å for the tunnels Ch2B-F, Ch2C, and the solvent channel Ch2S for generation of TCGnarrow and bottleneck radii of 2.0 Å for the tunnel Ch2B-F generating TCGwide (**Table S3**). These frames were then extracted and subject to five replicas of minimization, equilibration and followed by restrained NPT simulations as described for haloalkane dehalogenase.

## Results and discussion

### Tunnel networks formed in DhaA with different starting conformations and water models

Unsurprisingly, all restrained simulations of haloalkane dehalogenase showed good structural integrity with all three water models based on their root mean square deviation (RMSD) and root mean square fluctuation (RMSF) (**Figures S1 and S2)**. For all TCGs, RMSD values were consistently low, averaging between 0.45 Å and 0.5 Å. Additionally, domain movements across simulations using different models were similar, as seen by their RMSF. The fluctuations were primarily around 0.4 Å^2^ except for C- and N-terminal loops reaching up to 0.8 Å^2^.

The bottleneck radii of the P1 tunnel ensemble increased progressively along the predefined groups from TCG1 to TCG4 (**Figure 2**). Moreover, the tunnel radii observed in simulations starting from the same protein conformation followed a similar trend among all studied water models (**Figure S3**). This observation mostly aligns with the changes in the distance between the helices embedding the P1 tunnel, irrespective of the water model used in simulations (**Figure S4**). The water model also has a negligible effect on the composition of the bottleneck-forming residues (**Figure S5**), suggesting that the choice of water model did not markedly alter the location of the bottlenecks within the tunnels observed in those restrained simulations. Curiously, we have observed a systematic decrease in the bottleneck radii of all tunnels (**Figures 2 and S3**) compared to the radii observed in their initial starting structures (**Table S1**). Given the probe radius of 0.9 Å used for CAVER analyses, the reduced dimension of tunnels in TCG0, which was initiated from the structure with a bottleneck radius of 1 Å, became more evident when considering the almost halved frequency of detection of P1 tunnels in this TCG compared to TCGs1-4 for all water models (**Figure S6**). Considering other putative tunnels in the tunnel network, among many infrequent and narrow tunnels, two known auxiliary tunnels, P2 and P3, could be identified in simulations with all water models (**Figure S6**). However, the occurrences of these tunnels were less prevalent compared to the main P1 tunnel (**Figure S6)**. Similar observation was found by Klvana et al.^58^ where they observed the P2 tunnel occurring transiently and P3 tunnel was not present at all when investigating water transport through P1, P2 a,b,c and P3 tunnels. Moreover, these auxiliary tunnels also exhibited very narrow bottleneck regions, making them less viable for routine water transport (**Figure S7**), confirming the focus of the study on water transport via different well-defined geometries of the primary P1 tunnel.

**Figure 2:**
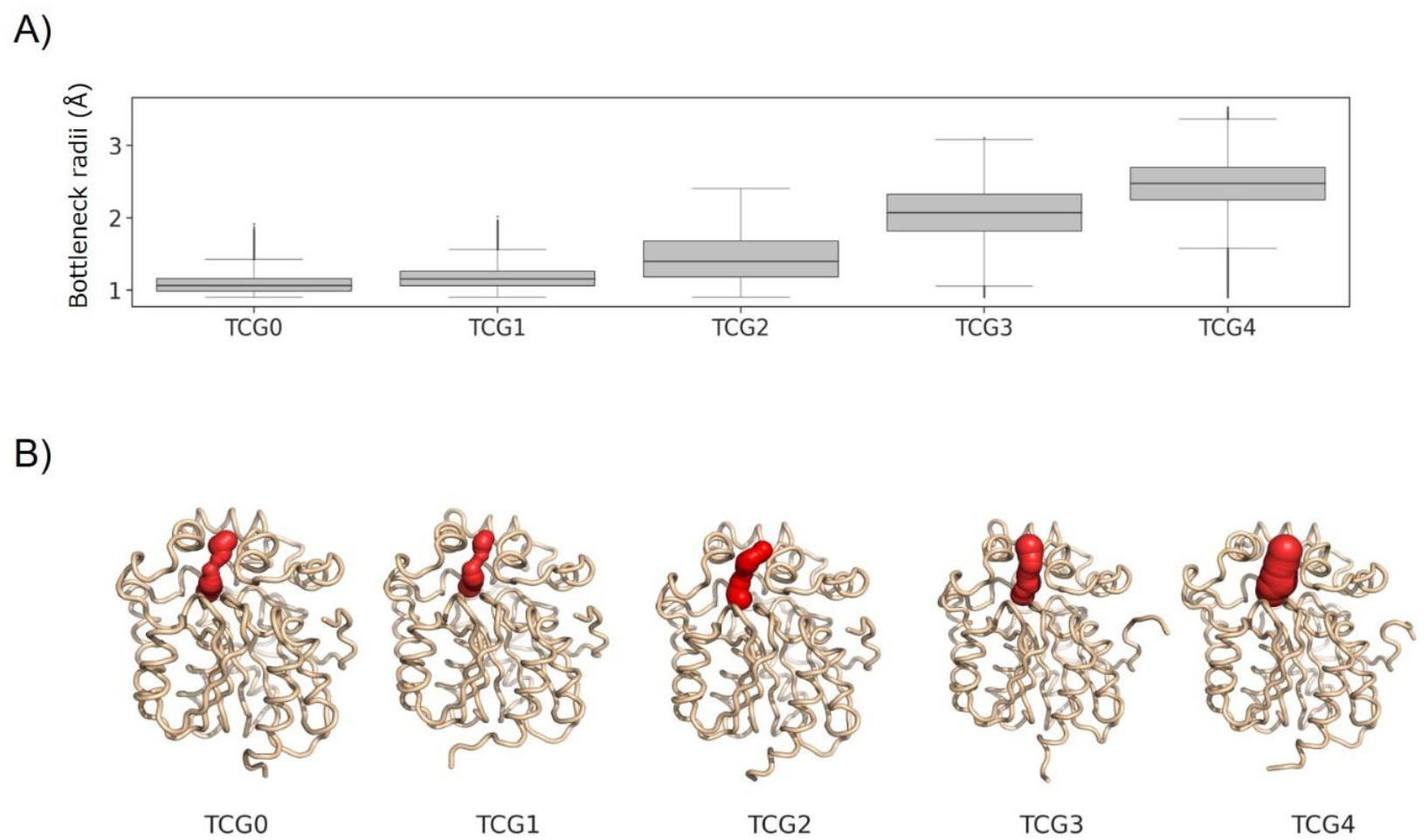
Comparison of P1 tunnel geometry sampled in DhaA simulations with different TCGs. (A) The average bottleneck radii of the P1 tunnel in different TCGs. For illustration, TCG0 and TCG1 exhibited sub-water radii (< Å), TSG2 matched the water dimension, while TSG3 and TSCG4 were widely open for water molecules. The data represent average±SD across 5 replicated simulations. (B) The representative structure of DhaA and P1 tunnel for TCG. The DhaA is visualized as ribbons, with the P1 tunnel shown with red spheres.

### Water transport through the P1 tunnel of distinct well-defined geometries is systematically different among water models

Upon analyzing the number of transported water molecules through P1 tunnels, we detected sparse water migration in only some of the simulations from TCG0 (**Figure S8**). This sparse data prevented us from quantitatively contrasting the water behavior within this TCG among the three studied water models. Such a rare water transport can be attributed to the small dimension of the P1 tunnel in TCG0, which was detected in only approximately 50% of the frames using a probe radius of 0.9 Å (**Figure S6**).

For all remaining TCGs, we have observed notable disparities among simulations using different water models in water transport via tunnels sharing identical bottleneck geometries and residue compositions (**Figure 3**). Approximately 2.5-fold larger amounts of water molecules (2.2-2.8; 95% confidence interval, CI) were transported in simulations with an open P1 tunnel (TCG2-4) using the TIP3P water model compared to OPC (**Figure 3A, B**). When we compared the transported TIP3P molecules to TIP4P-Ew ones, we observed smaller increase of about 2.0-fold (1.7-2.3; 95% CI). Overall, these observations applied to both the release of water molecules from the active site via the P1 tunnel and the entry of these molecules to the active site via the same route (**Figure S8**). Given the rare nature of transport events through rather narrow tunnels in TCG1 (**Figures S8 and 2A**), we have not observed statistically significant differences among the models in this TCG.

**Figure 3:**
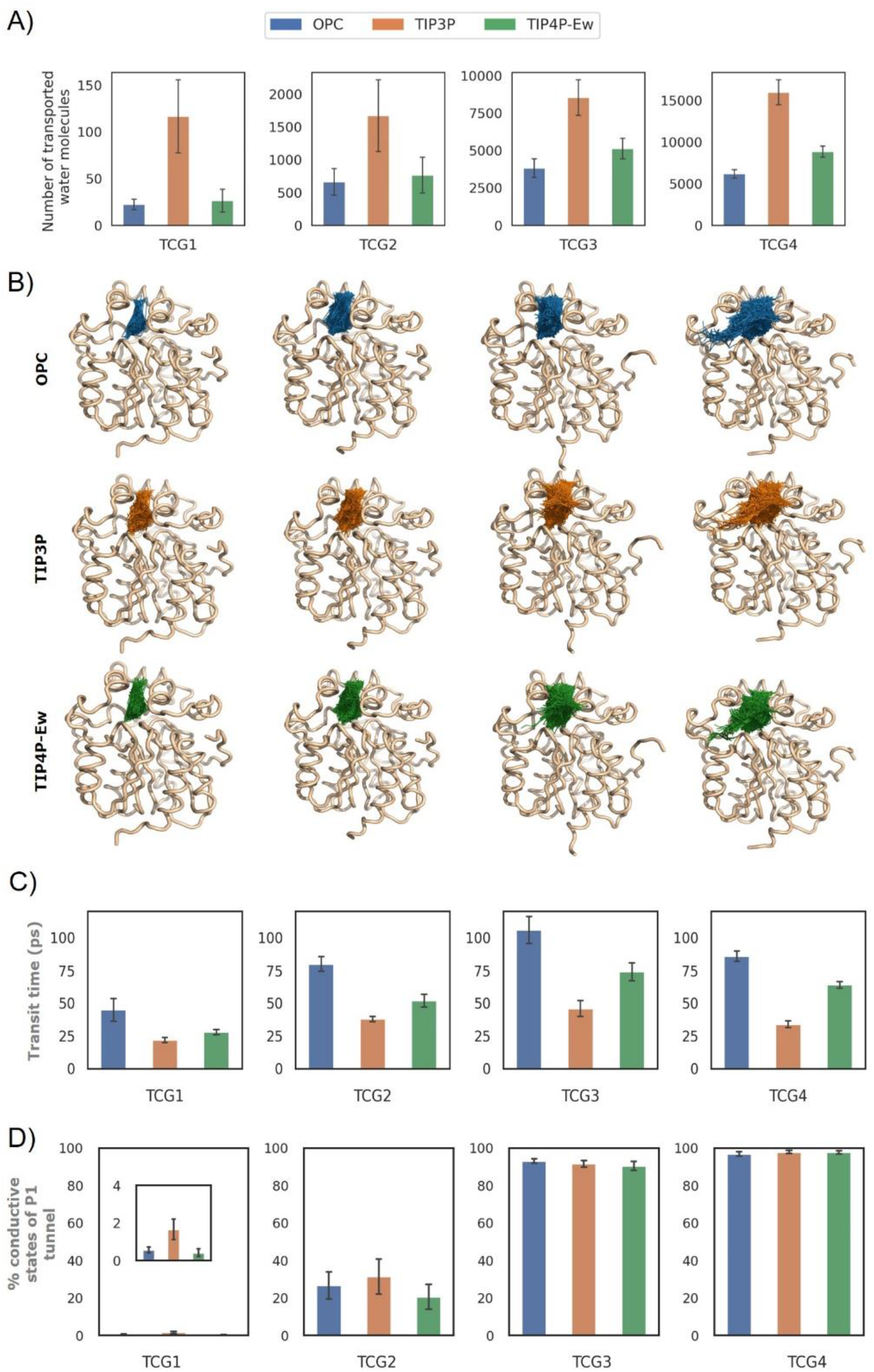
Impact of different water models on the transport of water molecules through P1 tunnels of DhaA in different TCGs. (A) The total number of water molecules transported through P1 tunnels in different TCGs and different models. (B) The representative protein structure with traces of water molecules migrating via the P1 tunnel. (C) Median transit time for water molecules migration across P1 tunnel (D) Percentage of conductive states adopted by P1 tunnel for water molecules. The data in A), C), and D) represent average±SEM across 5 replicated simulations.

To understand the origin of these differences, we first investigated the duration of water transit between the bulk solvent and the active site of DhaA with different models (**Figures 3C and S9**). We observed that the TIP3P water molecules exhibited approximately 2.3-fold (2.1-2.5;95% CI) faster movement across the tunnels than OPC, while TIP4P-Ew water molecules migrated at about 1.6-fold (1.4-1.7;95% CI) of TIP3P mean transit times. Additionally, we observed a tendency towards longer transit times with more open tunnel geometries. This observation in TCG3-4 could be a result of water forming static networks which can increase the residence time of water in the tunnel,^9^ and can form clusters of waters during migration through the tunnel. ^59^ Although TIP3P water molecules were the fastest to migrate through the tunnels, this difference might not be sufficient to fully explain the 2.5 times larger number of transported water molecules with this model observed in TCG2-4 (**Figure 3A**).

Hence, we have analyzed the number of water molecules that were transported through the P1 tunnel at the same time (**Figure S10**). As expected, the number of concurrently transported molecules reflected the dimension of the P1 tunnel and exhibited about a 3.8-fold increase from TCG1 to TCG4 (3.3-4.3; 95% CI), transforming from essentially a single molecule transport in TCG1&2 to simultaneous transport of 3-4 water molecules in TCG3&4. Interestingly, only tunnels from TCG3&4 enabled considerable bi-directional migration of water molecules, while the tunnels from TCG1&2 allowed mainly unidirectional movements. Often in narrow regions like in TCG1-2, water movement can be restricted by side chain conformation like found in aquaporin^60,61^ and in other studies waters was found to migrate in a single chain to the cavity, ^62,63^ which still could be affected by energy barriers during the migration.^64^ However, we have not observed any notable differences among the water models in the parallel nature of the migration of water molecules, which is in agreement with the consistent tunnel geometries observed in the same TCGs among the water models due to applied restraints. Finally, we have evaluated the frequency with which the P1 tunnel was observed in states conducting at least one water molecule (**Figure 3D**), which indicated that TIP3P water molecules might be able to utilize narrow regions of the tunnels in TCG1 more often than the other two studied models, unlike in TCG2-4. The wider tunnel geometries in TCG4&5 enabled almost continuous utilization of the P1 tunnel for water migration across all simulations.

### Impact of water models on water transport in other enzymatic systems

Similarly to dehalogenase, restrained simulations of AldO and Cyt P450 exhibited excellent structural integrity with all tested water models (**Figures S11-S14**). For AldO, we have observed the investigated tunnel T1 as the primary one in both TCGnarrow and TCGwide (**Figure S15**). T1 in AldO defines the entrance to the catalytic pocket for the substrate^65^ and is considered as the primary tunnel for water transportation as the other narrower funnels (like T2) facilitate only the diffusion of the molecular oxygen to the active site. ^66^ In the case of Cyt P450, the merged Ch2F-B tunnels mainly serve for substrate access, the Ch2S tunnel, which is called the solvent channel, may play a role in a metabolite egress channel, and the Ch2C tunnel contributes to substrate access.^67–69^ They were the most observed tunnels in TCGnarrow and TCGwide regardless of the water models used (**Figure S16**).

Considering the sampled dimensions of these tunnels, TCGnarrow bottleneck radii increased in comparison to the selected geometries for T1 in AldO as well as Ch2F-B and ChS in Cyt P450, while TCGwide bottleneck radii of T1 in AldO as well as Ch2F-B and ChS in Cyt P450 were slightly decreased in comparison to initially selected structures (**Figure 4A and Tables S2 and S3**). In contrast, bottleneck radii of the Ch2C tunnel of CytP450 remained very narrow in both TCGs. The bottleneck radii of the T1 tunnel of AldO in both TCGs, as well as Ch2F-B and ChS tunnels of Cyt P450 in TCGwide, were suitably opened for water molecules similarly to TCG3&4 of DhaA. Ch2F-B and ChS tunnels of Cyt P450 in TCGnarrow matched the water dimension like TCG2 of DhaA (**Figures 4A and 2A**). Finally, the ChC tunnel of Cyt P450 exhibited sub-water radii in both TCGs similarly to TCG0&1 of DhaA.

**Figure 4:**
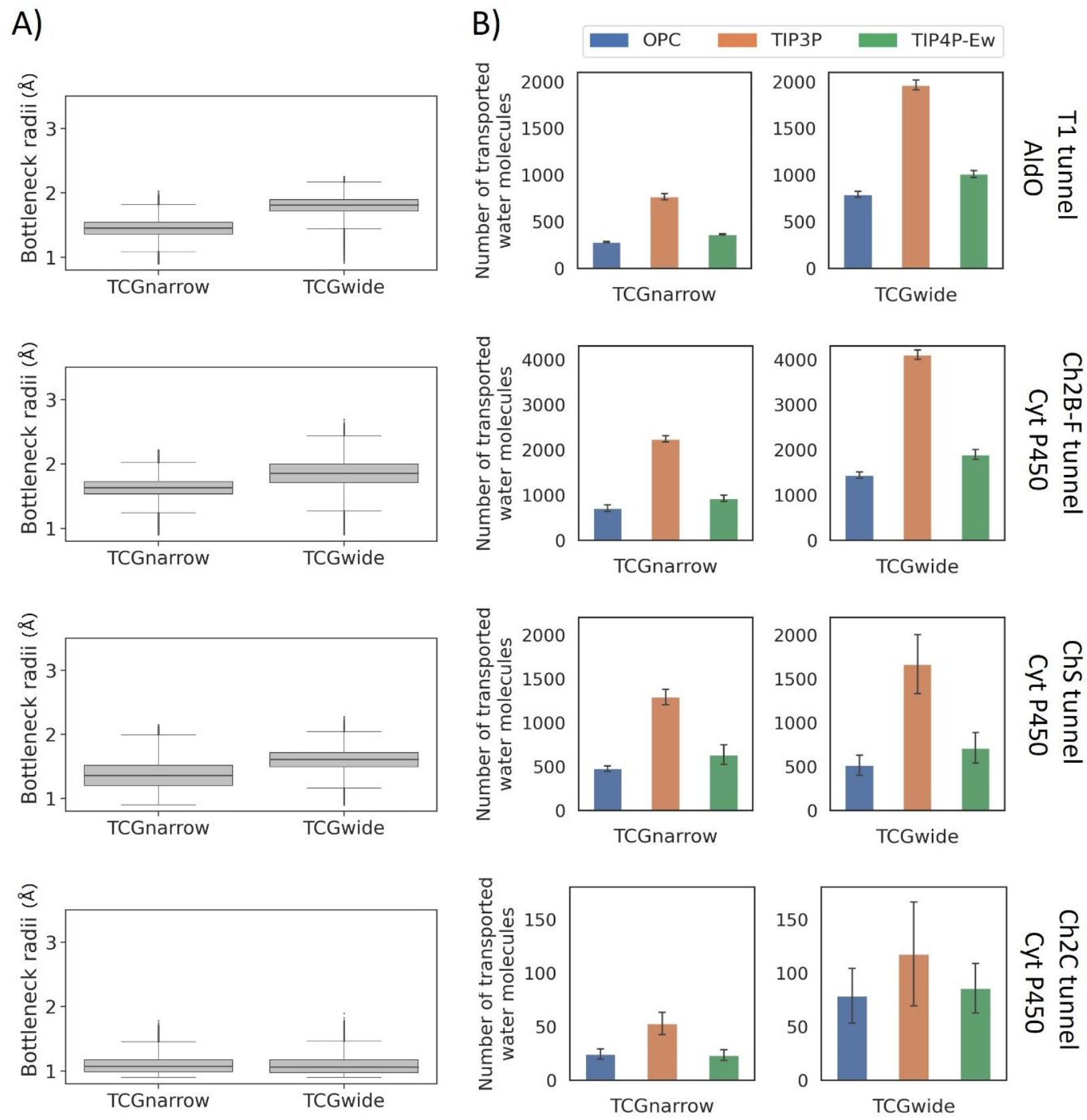
Impact of different water models on the transport of water molecules through tunnels of AldO and Cyt P450 in two defined TCGs. (A) The average bottleneck radii of the investigated tunnels in the two TCGs. (B) The total number of water molecules transported through investigated tunnels in two TCGs and different models. The data represent average±SEM across 5 replicated simulations.

The water transport across the tunnels in both enzymes exhibited similar trends as observed with haloalkane dehalogenase DhaA (**Figure 4B and Figures S19-S30**). The amount of transported water molecules was still about 2.5-fold increased with TIP3P compared to OPC (95% CI of 2.4-2.8 and 2.4-2.9 for AldO and Cyt P450, respectively) and about twice larger over TIP4P-Ew (95% CI of 1.9-2.2 and 2.0-2.3 for AldO and Cyt P450, respectively). These differences among the three studied water models were consistent with the differences in the transit times of water molecules through the investigated tunnels (**Figures S19-S22**). We have not observed any significant differences among the number of water molecules that were transported in parallel (**Figures S23-S26**) nor the frequency of adopting conductive states by the investigated tunnels among different water models (**Figures S27-S30**). While the overall number of transported water molecules generally reflected the dimension of the utilized tunnel similarly to DhaA, we have noted interesting behavior with the Ch2C tunnel of Cyp P450 which transported more than twice the amount of water molecules in TCGwide than in TCGnarrow despite having equivalent bottleneck radii in both TCGs (**Figure 4**). This observation could be traced to the notably increased utilization of this relatively narrow ChC tunnel for the flow of water molecules in TCGwide (**Figure S30**), possibly due to the increased water transport via the other two more opened tunnels (Ch2B-F and ChS). Overall, our data with AldO and Cyt P450 confirmed systematic differences in the utilization of tunnels when using different water models irrespective of their geometry, residue composition, localization in the protein, and functional roles.

## Conclusions

In this study, we performed simulations with different fixed geometries of tunnels, enabling us to investigate the effects of different water models on water transport through well-controlled tunnels. Such a setup allowed us to assign the differences in the number and behavior of transported water molecules through the same tunnels in different simulations to the differences in the physicochemical properties of the water molecules used in the simulations. TIP3P is the most popular water model with researchers, and it is the most widely used water. In the context of water transport through tunnels, the diffusivity of the water model used will play a key role in determining the utilization of a tunnel. We observed in all situations that TIP3P waters migrate faster and even enter the narrow tunnel regions more willingly than with the other two analyzed water models. Such preference might be due to the increased ability of TIP3P water molecules to escape the bulk solvent, given by their much higher self-diffusion coefficient (5.72 ± 0.04 ×10^−5^ cm^2^/s) when compared to OPC (2.27 ± 0.02 ×10^−5^ cm^2^/s), TIP4P-Ew (2.54 ± 0.01 ×10^−5^ cm^2^/s), and experimental observations (2.30 ×10^−5^ cm^2^/s).^70,71^ We confirmed our observations of this phenomenon in two other different enzymes with buried active sites and different topologies of tunnel networks, suggesting we can extrapolate our findings to most of the enzymes that utilize well-defined tunnels to exchange small molecules between their buried active sites and the bulk solvent. Considering the importance of water molecules’ availability during different stages of the catalytic cycle in many enzymes and the role of water molecules in unbinding the reaction products and inhibitors, the choice of a proper water model will be critical for simulating complex enzyme reactions,^72–75^ rationally designing enzymes,^76,77^ and accurately predicting the drug residence times. ^78–80^

## Acknowledgments

This work was supported by the National Science Centre, Poland (grant no. 2017/26/E/NZ1/00548 to J.B.). A.S.T was the recipient of POWR.03.02.00-00-I006/17 scholarship and grants of the dean in faculty of Biology, Adam Mickiewicz University in Poznan GDWB05/2020. The simulations and analyses of DhaA were performed in the Poznan supercomputing center. Polands high-performance Infrastructure PLGrid (HPC Centers: ACK Cyfronet AGH:ares) is gratefully acknowledged for providing computer facilities and support for analyses of AldO and Cyt P450.

## Author contributions

Conceptualization: J.B., A.G.; Data curation: A.S.T (DhaA), K.S. (AldO), G.T. (Cyt P450); Formal analysis: A.S.T (DhaA), K.S. (AldO), G.T. (Cyt P450); Funding acquisition: J.B., AG; Investigation: A.S.T. (DhaA), KSz (AldO), GT (Cyt P450); Methodology: A.S.T; Project administration: J.B., A.G; Resources: J.B., K.M.; Software: J.B., K.M.; Supervision: J.B., A.G. Validation: J.B., A.G.; Writing – original draft: A.S.T., KSz (AldO), GT (Cyt P450); Writing – review & editing: J.B., K.M., A.G.

