## Supporting information for "Water migration through enzyme tunnels is sensitive to choice of explicit water model"

**Table S1: Details on the initial structure for restrained simulation of haloalkane dehalogenase DhaA.** The initial structure for simulation was obtained from a 5 µs HTMD (High-Throughput Molecular Dynamics) simulation described at https://doi.org/10.1101/2023.05.24.542065. From this simulation, 15 frames were selected based on the bottleneck radii of the P1 tunnel.

| Group | HTMD simulation ID | Frame | Bottleneck radii [Å] |
|---|---|---|---|
| | e6s1_e5s1p0f1660 | 7090 | **1.00** |
| | e10s1_e9s3p0f1600 | 6486 | **1.00** |
| **TCG0** | e1s5_conf1 | 2510 | **1.00** |
| | e3s3_e2s2p0f1430 | 5456 | **1.00** |
| | e2s5_e1s2p0f1850 | 7896 | **1.00** |
| | e9s1_e8s2p0f8840 | 909 | **1.40** |
| | e4s3_e3s5p0f3540 | 9986 | **1.40** |
| **TCG1** | e1s3_conf1 | 6703 | **1.40** |
| | e8s2_e7s5p0f8080 | 8191 | **1.40** |
| | e8s4_e7s5p0f9500 | 3824 | **1.40** |
| | e1s3_conf1 | 7833 | **1.80** |
| | e9s5_e8s1p0f4660 | 7810 | **1.80** |
| **TCG2** | e1s3_conf1 | 4855 | **1.80** |
| | e5s4_e3s5p0f1180 | 8347 | **1.80** |
| | e5s5_e1s2p0f2860 | 4468 | **1.80** |
| | e5s4_e3s5p0f1180 | 9434 | **2.49** |
| | e9s5_e8s1p0f4660 | 6776 | **2.49** |
| **TCG3** | e5s4_e3s5p0f1180 | 8137 | **2.48** |
| | e9s5_e8s1p0f4660 | 6178 | **2.47** |
| | e6s2_e5s1p0f620 | 4927 | **2.47** |
| | e5s4_e3s5p0f1180 | 7716 | **3.08** |
| | e5s4_e3s5p0f1180 | 7722 | **3.07** |
| **TCG4** | e6s2_e5s1p0f620 | 4923 | **2.90** |
| | e5s4_e3s5p0f1180 | 8097 | **2.89** |
| | e6s2_e5s1p0f620 | 4925 | **2.86** |

**Table S2: Details on the initial structure for restrained simulation of alditol oxidase.** Frames were selected based on the bottleneck radii of the T1 tunnel.

| Group | Frame | Bottleneck radii [Å] |
|---|---|---|
| **TCGnarrow** | 3743 | 1.00 |
| **TCGwide** | 23729 | 2.00 |

**Table S3: Details on the initial structure for restrained simulation of cytochrome P450.** Frames were selected based on the bottleneck radii of the Ch2B-F, ChS and Ch2C tunnels.

| | | | Bottleneck radii [Å] | |
|---|---|---|---|---|
| Group | Frame | Ch2B-F tunnel | ChS tunnel | Ch2C tunnel |
| TCGnarrow | 24648 | 1.14 | 1.14 | 1.14 |
| TCGwide | 10348 | 2.00 | 2.00 | 2.00 |

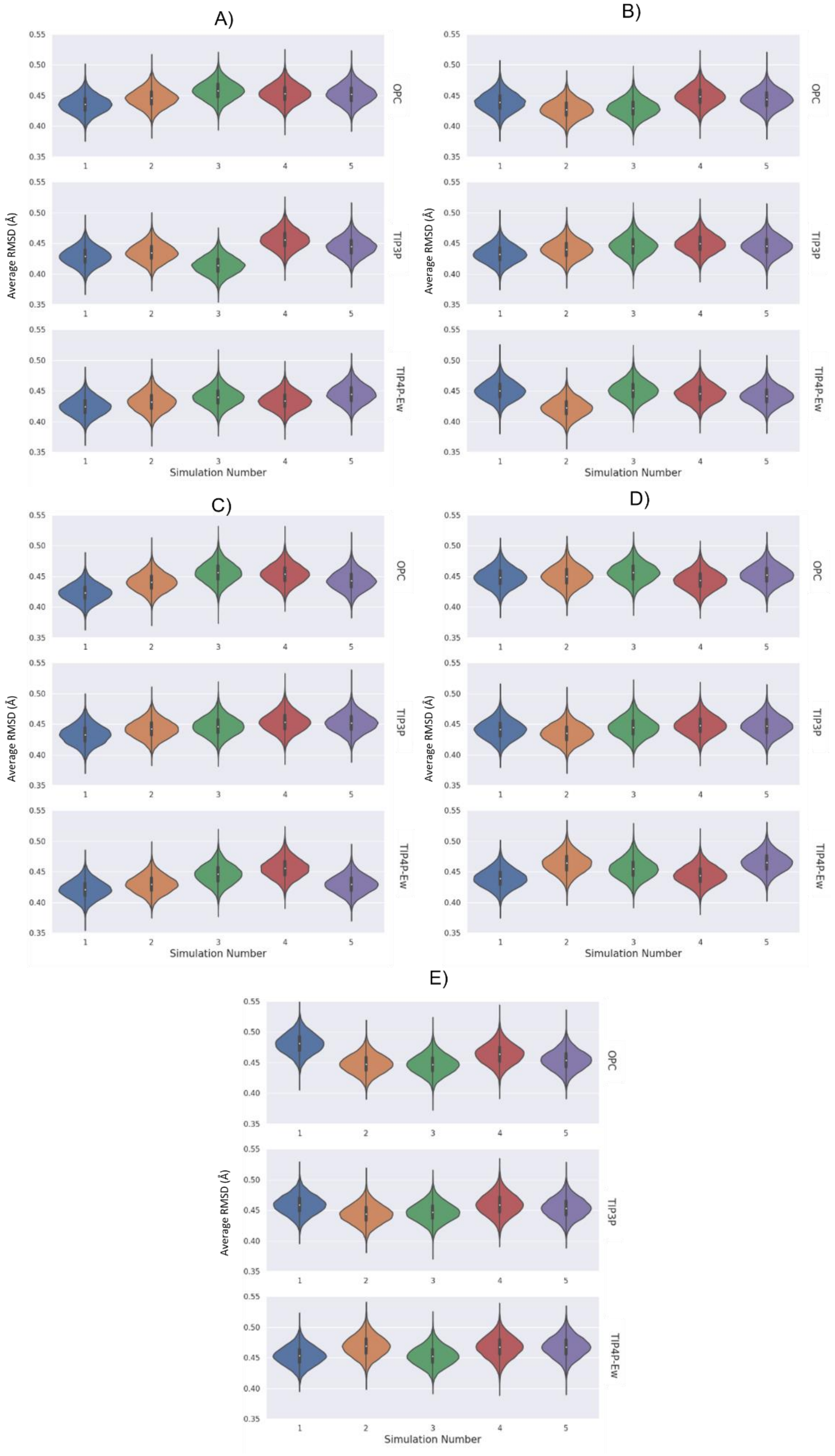

**Figure S1 Root mean squared deviations (RMSD) of backbone heavy atoms of haloalkane dehalogenase DhaA for TCG0(A), TCG1(B), TCG2(C), TCG3(D) and TCG4(E).**

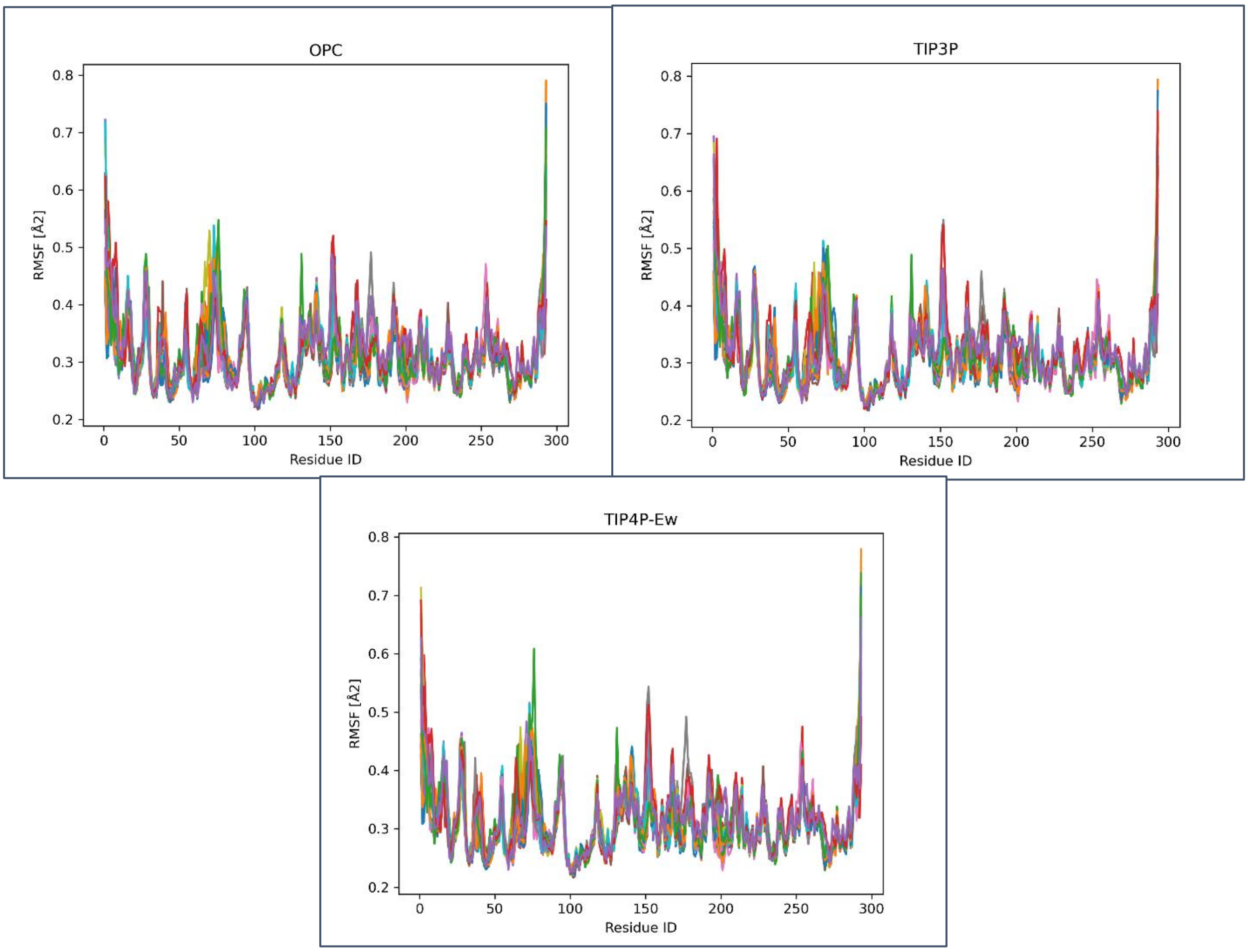

**Figure S2: Root Mean Square Fluctuation (RMSF) of haloalkane dehalogenase DhaA with studied water models.** The RMSF values were calculated for backbone heavy atoms of each residue in the protein to assess the flexibility and dynamic behavior of the enzyme under distinct solvation conditions.

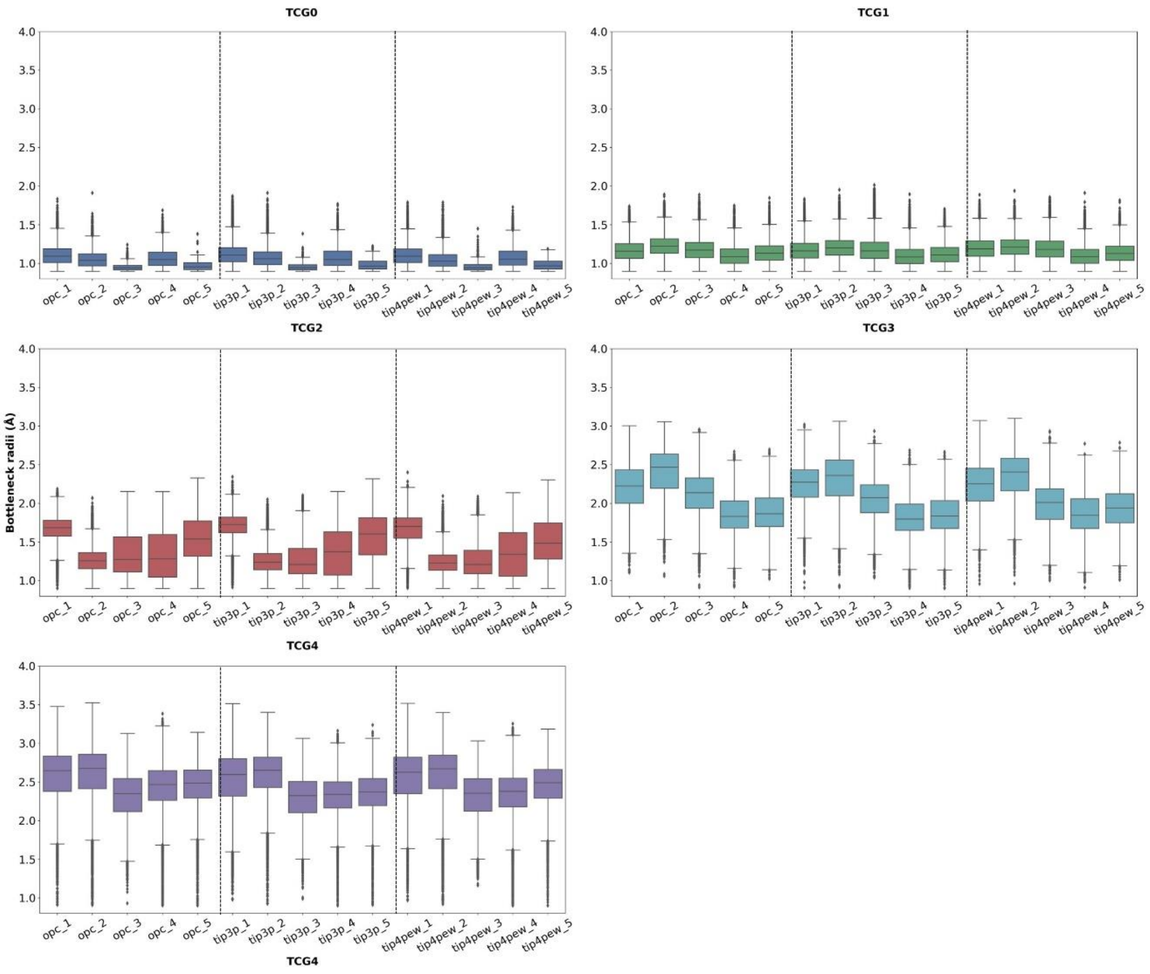

**Figure S3: The average bottleneck radii of P1 tunnel in different simulations of haloalkane dehalogenase DhaA**. For each TCG, the first box represents the average bottleneck observed in individual restrained simulations. The data repersent average±SD across 5 replicated simulations.

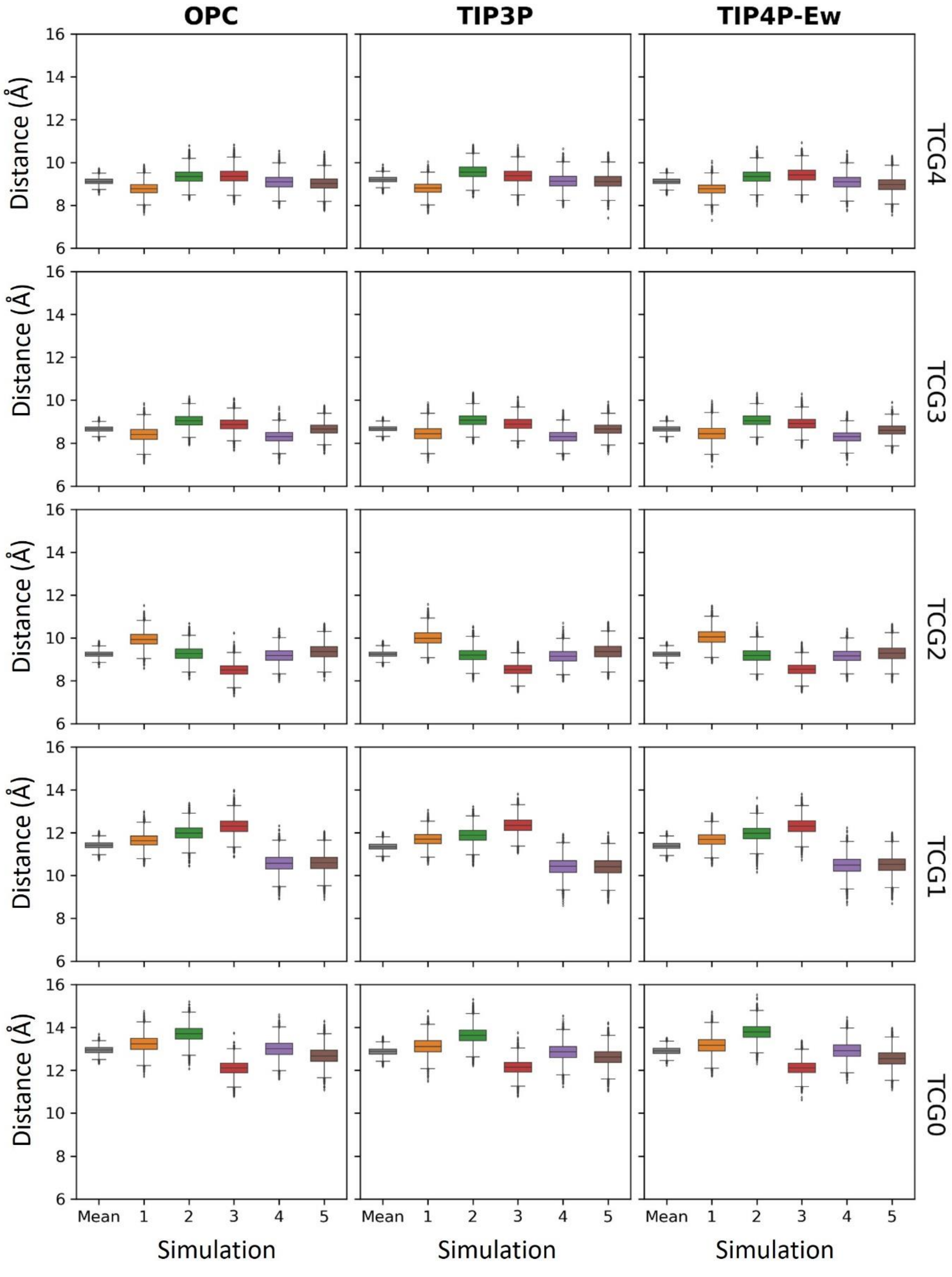

**Figure S4: Helix-helix distance at mouth of P1 tunnel of haloalkane dehalogenase DhaA measured between CA atoms of F144 and K175.** The data repersent average±SD across 5 replicated simulations.

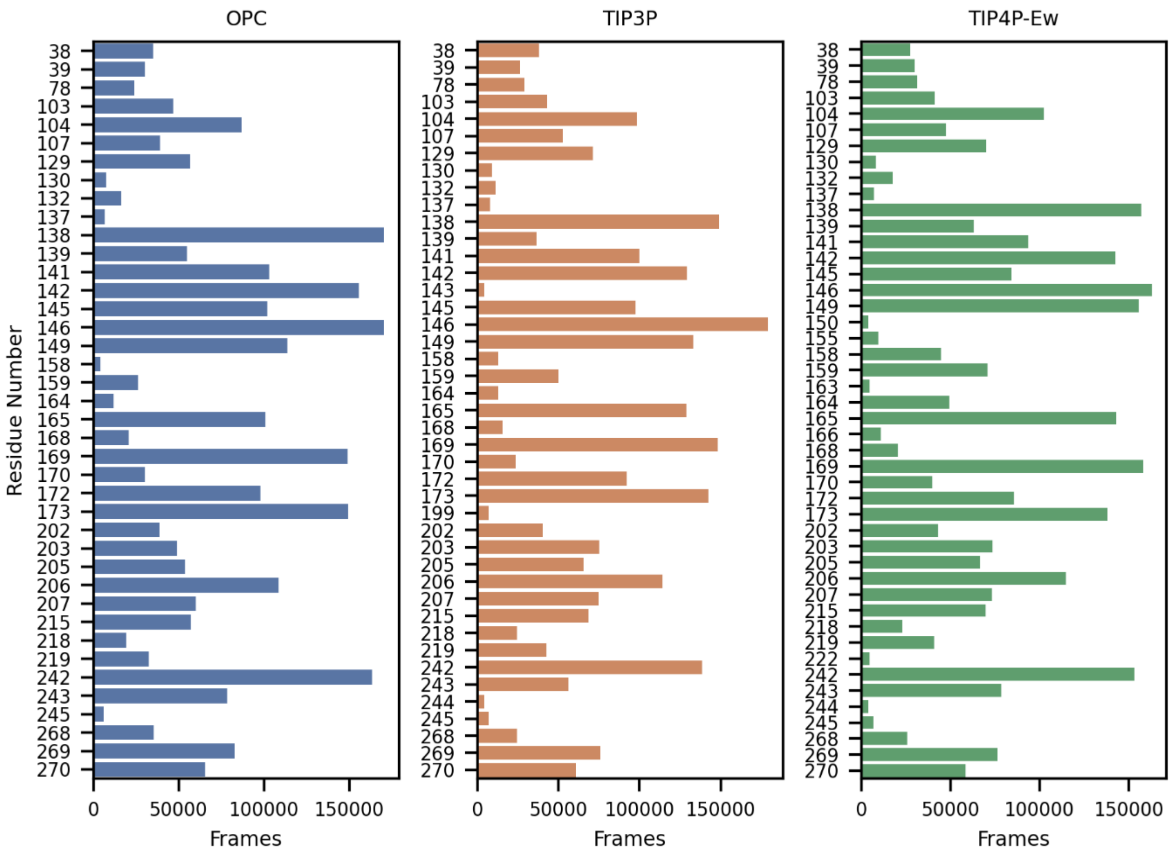

**Figure S5: Residues forming the bottlenecks of P1 tunnel of haloalkane dehalogenase DhaA during the simulation.**

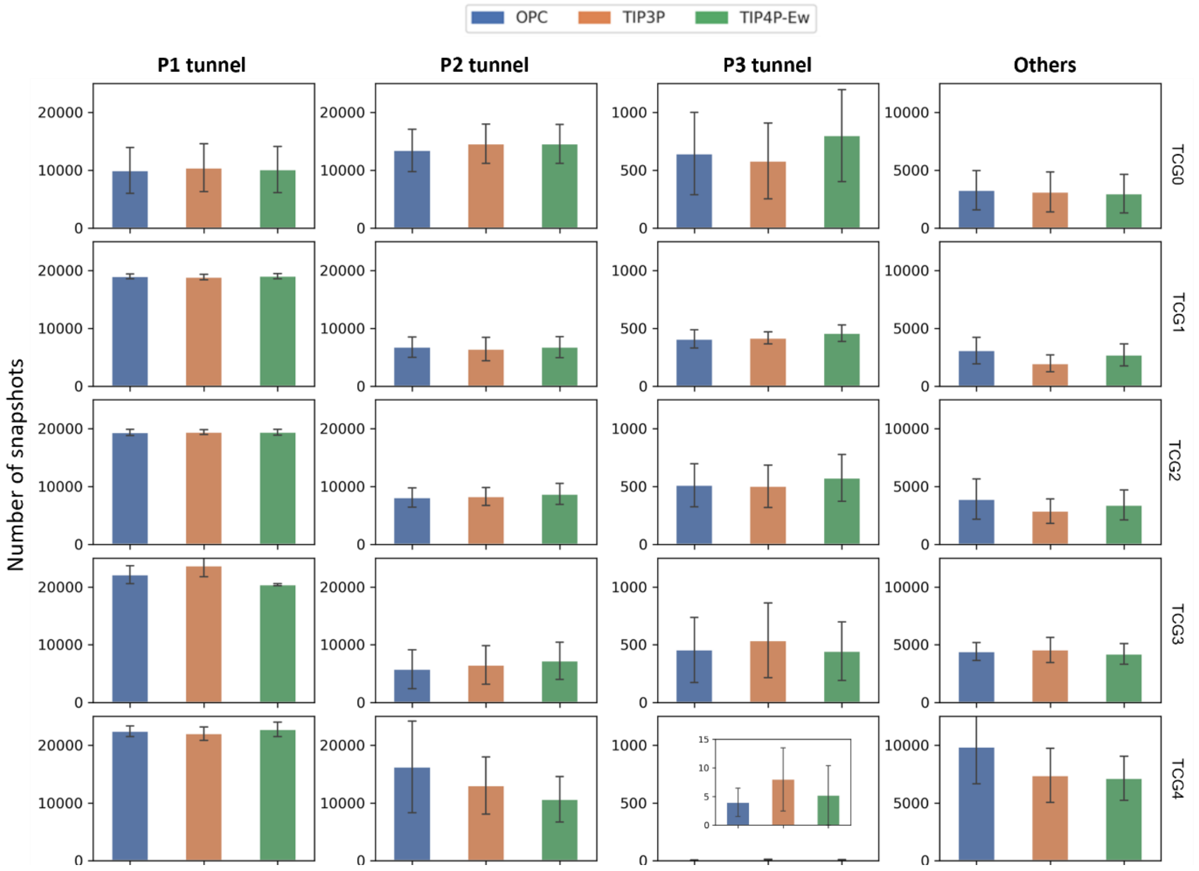

**Figure S6: Number of tunnels detected in simulations of haloalkane dehalogenase DhaA.** The data represent average±SEM across 5 replicated simulations.

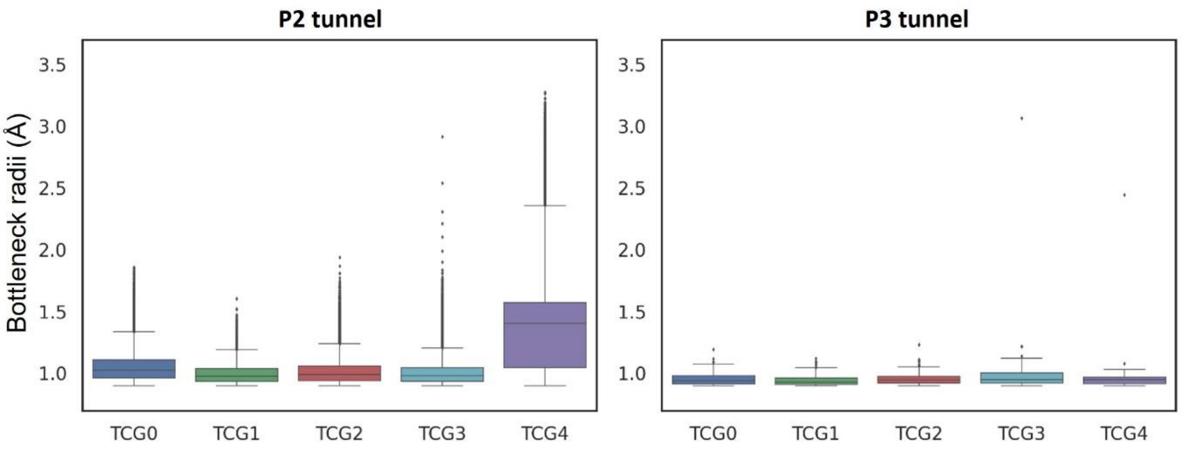

**Figure S7: Average Bottleneck radii of P2 and P3 tunnels of haloalkane dehalogenase DhaA in different TCGs.** The data represent average±SD across 5 replicated simulations.

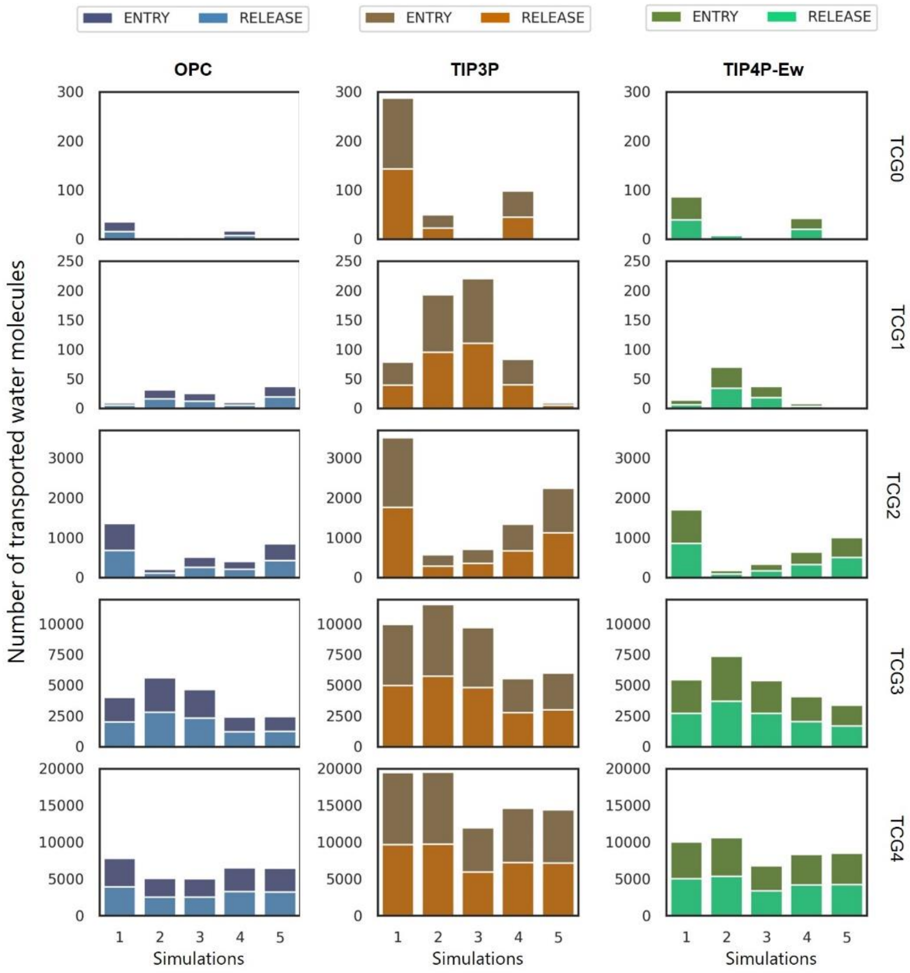

**Figure S8: Use of P1 tunnel in individual TCGs of haloalkane dehalogenase DhaA for entry of water molecules to the active site and their release to bulk with three studied water models.**

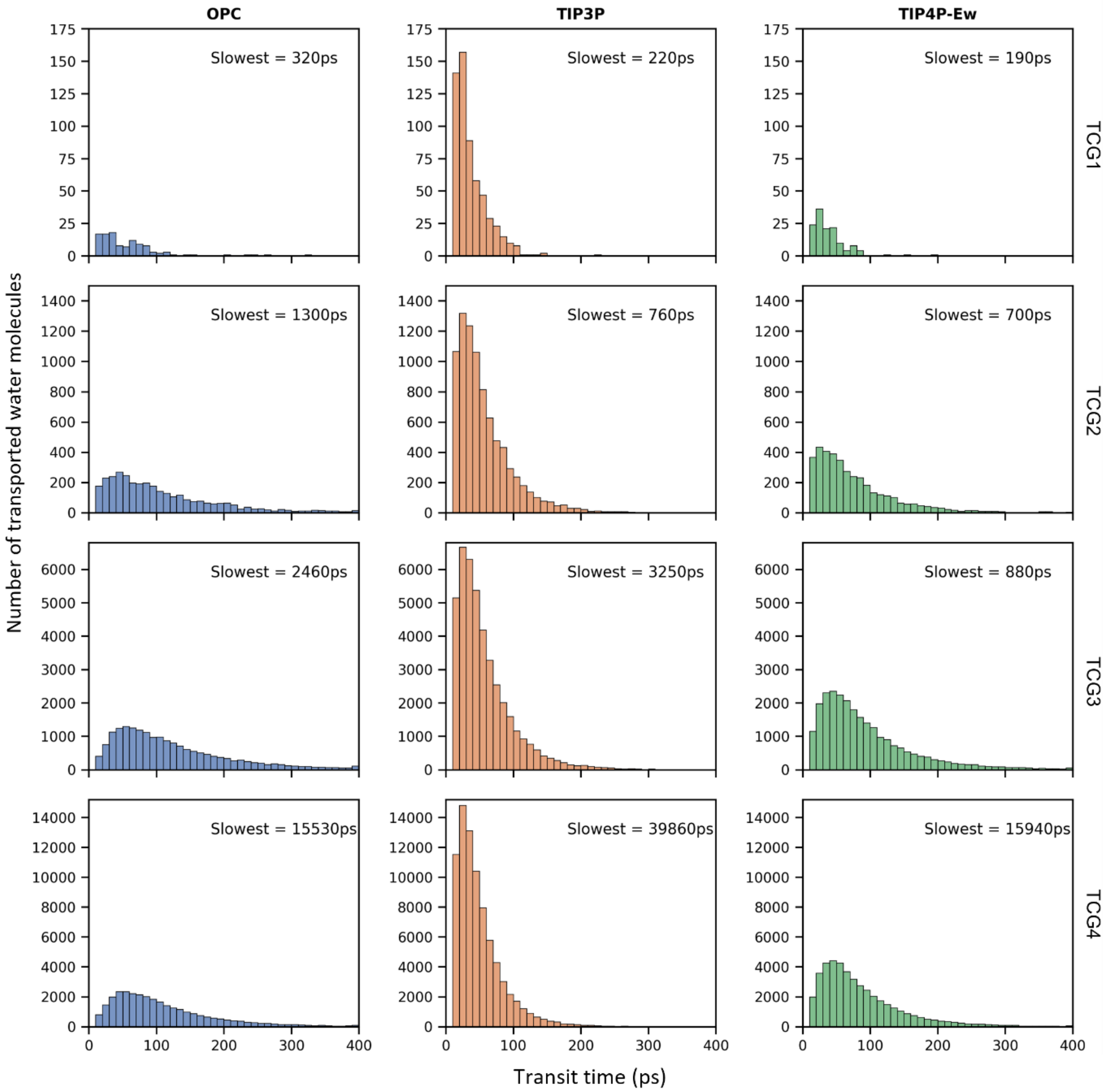

**Figure S9: Distribution of transit time of migrating water molecules through P1 tunnel of haloalkane dehalogenase DhaA.**

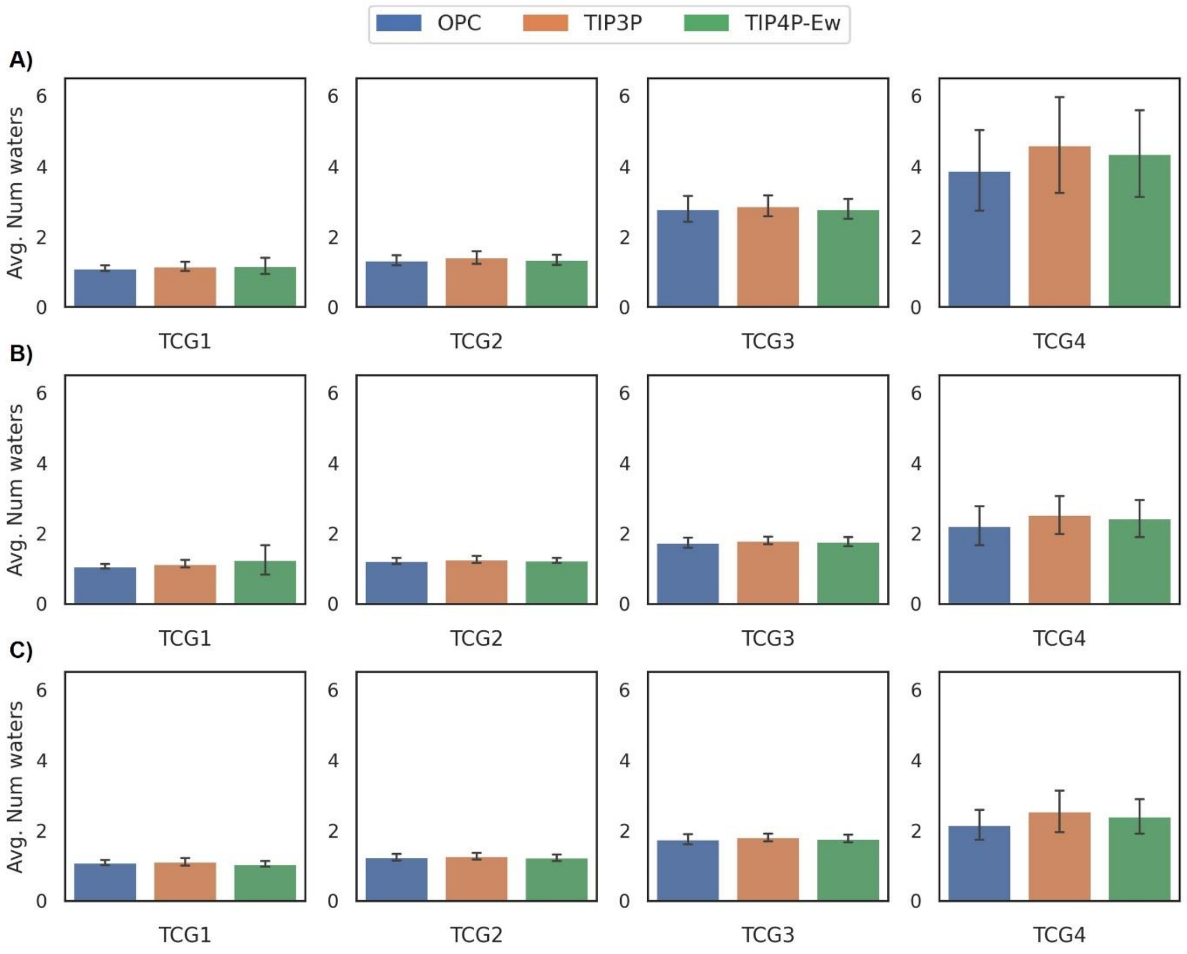

**Figure S10: Number of waters molecules concurrently transported via P1 tunnel of haloalkane dehalogenase DhaA.** The number of concurrent water molecules regardless the direction of their movement (A), entering the active site (B), and releasing to the bulk solvent (C). The data represent average±SEM across 5 replicated simulations.

A)

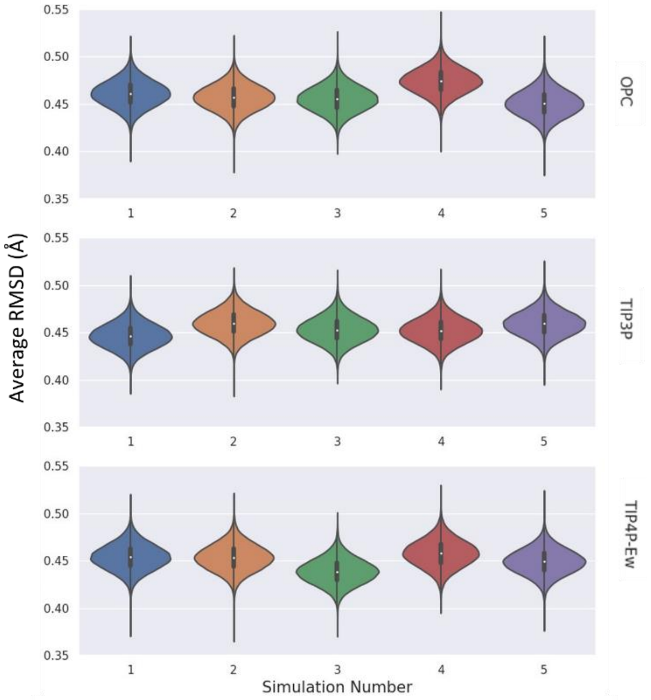

B)

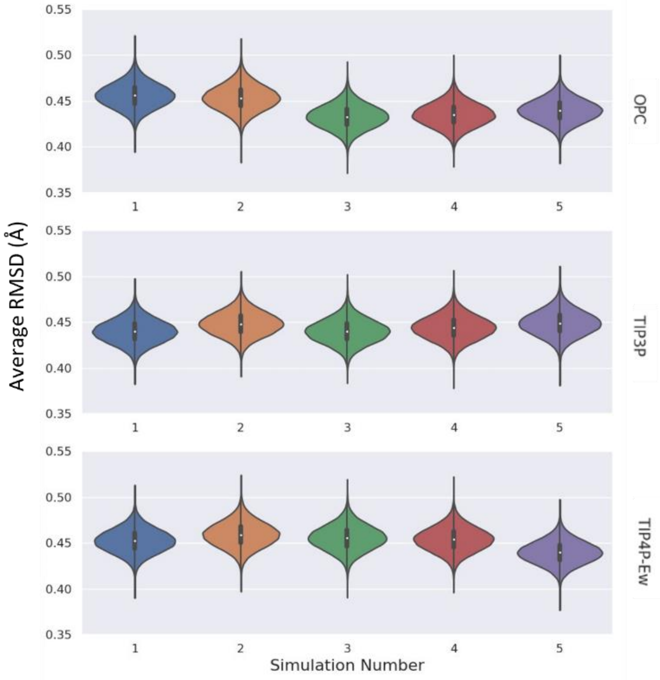

**Figure S11: Root mean squared deviations (RMSD) of backbone heavy atoms of alditol oxidase for TCGnarrow (A), and TCGwide (B).**

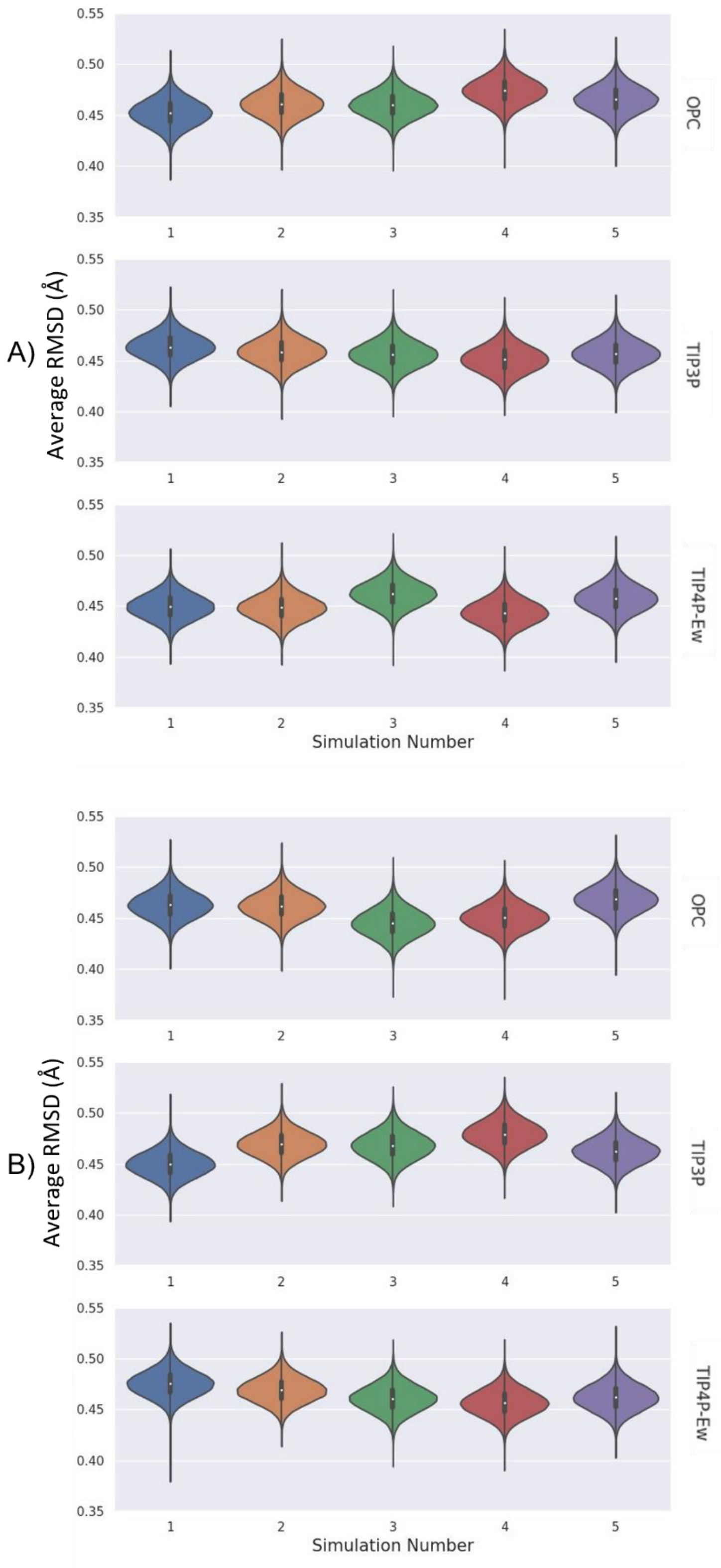

**Figure S12: Root mean squared deviations (RMSD) of backbone heavy atoms of cytochrome P450 for TCGnarrow (A), and TCGwide (B).**

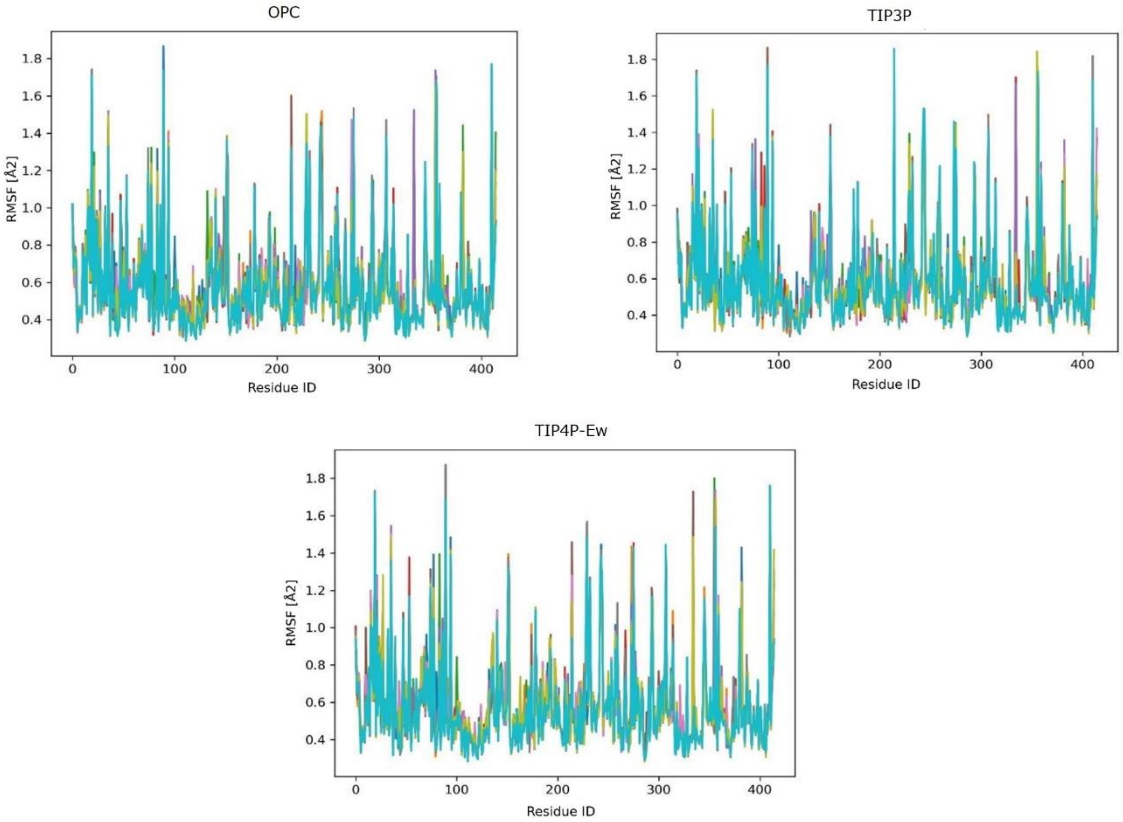

**Figure S13: Root Mean Square Fluctuation (RMSF) of alditol oxidase with studied water models.** The RMSF values were calculated for backbone heavy atoms of each residue in the protein to assess the flexibility and dynamic behavior of the enzyme under distinct solvation conditions.

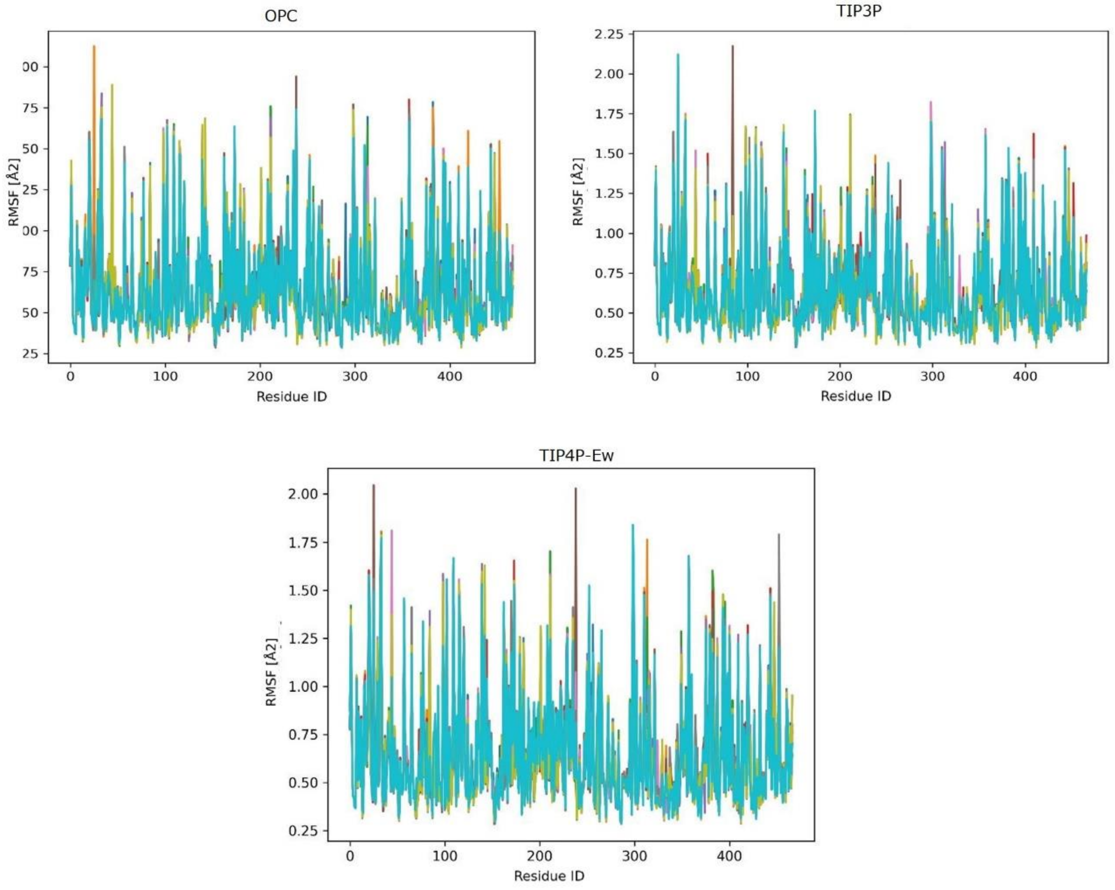

**Figure S14: Root Mean Square Fluctuation (RMSF) of cytochrome P450 with studied water models.** The RMSF values were calculated for backbone heavy atoms of each residue in the protein to assess the flexibility and dynamic behavior of the enzyme under distinct solvation conditions.

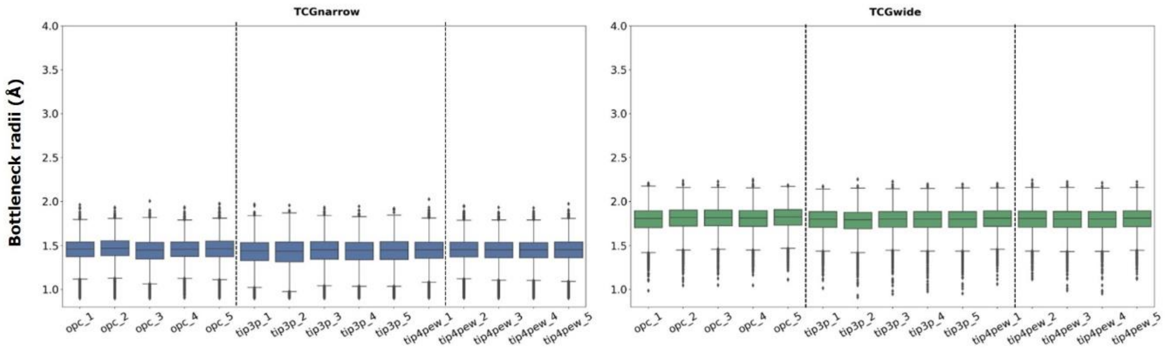

**Figure S15: The average bottleneck radii of T1 tunnel in different simulations of alditol oxidase**. For each TCG (narrow, wide), the boxes represents the average bottleneck radii observed in individual simulations. The data repersent average±SDM across 5 replicated simulations.

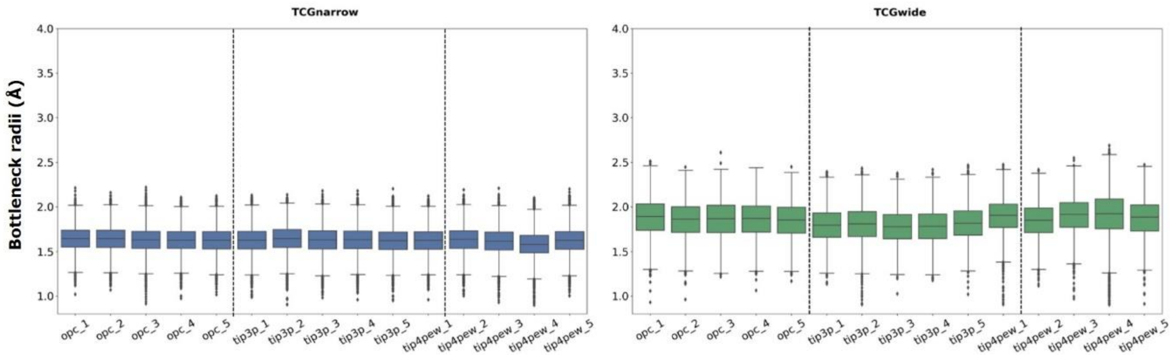

**Figure S16: The average bottleneck radii of Ch2B-F tunnel in different simulations of cytochrome P450**. For each TCG (narrow, wide), the boxes represents the average bottleneck radii observed in individual simulations. The data repersent average±SD across 5 replicated simulations.

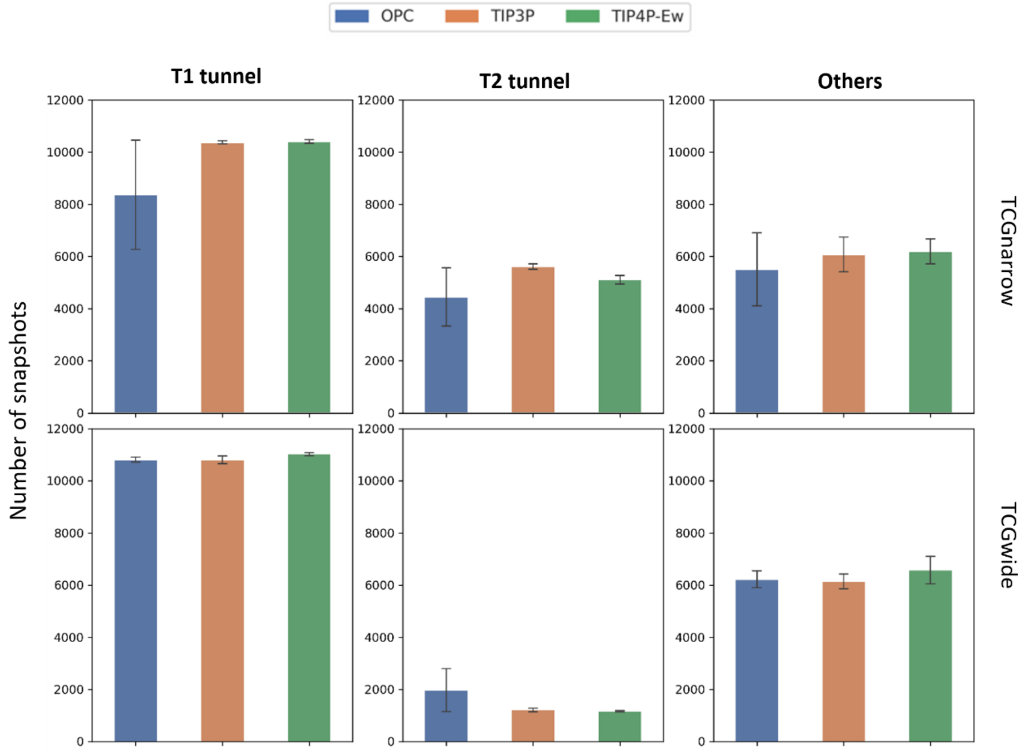

**Figure S17: Number of tunnels detected in simulations of alditol oxidase with narrow and wide tunnel groups.** The data repersent average±SEM across 5 replicated simulations.

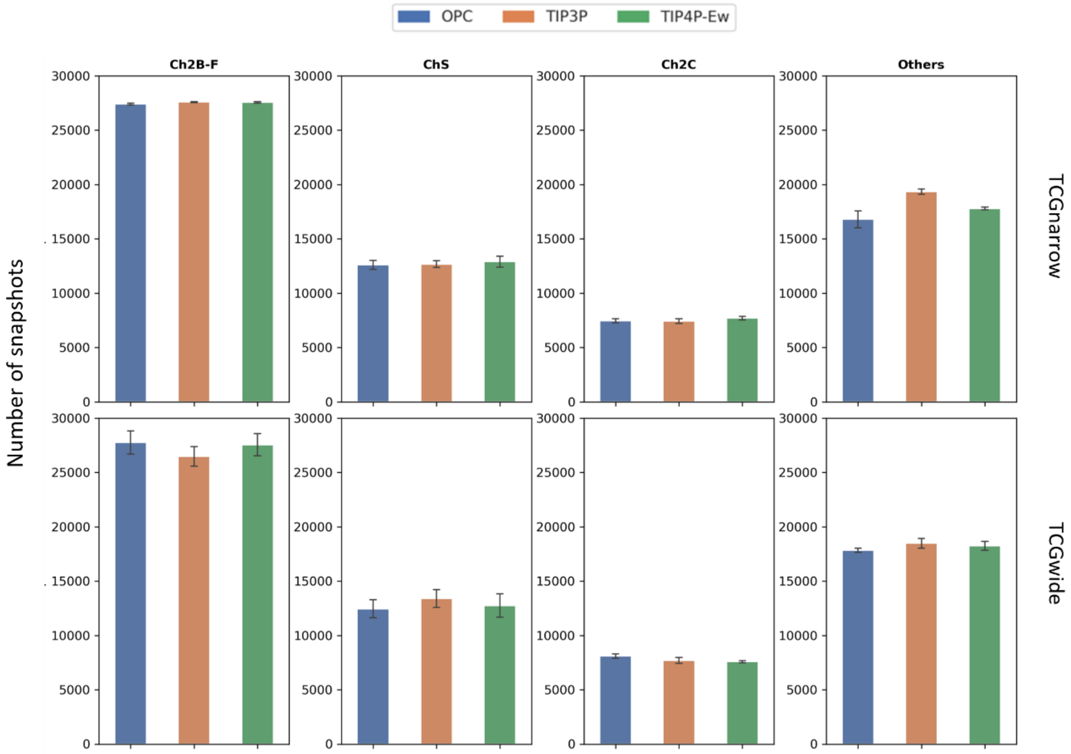

**Figure S18: Number of tunnels detected in simulations of cytochrome P450 with narrow and wide tunnel groups.** The data repersent average±SEM across 5 replicated simulations.

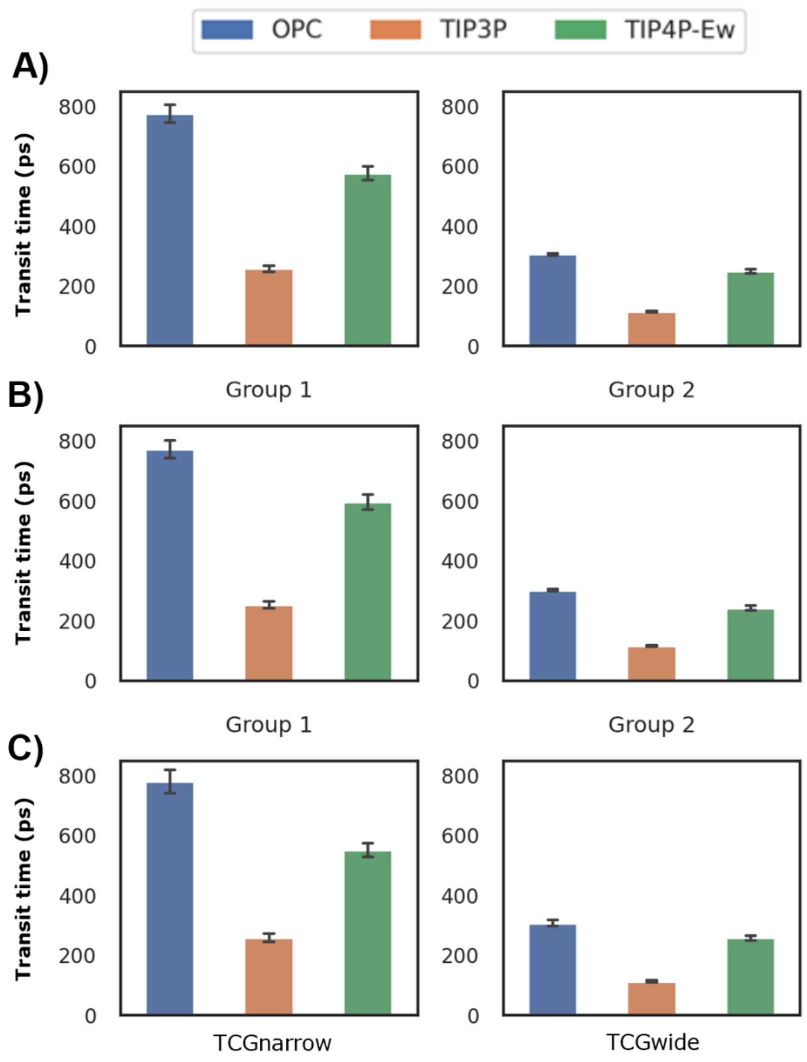

**Figure S19: Median transit time for water molecules migration across T1 tunnel for alditol oxidase.** Transit time regardless of the direction of water movement (A), water entering the active site (B), and water releasing to the bulk solvent (C). The data represent average±SEM across 5 replicated simulations.

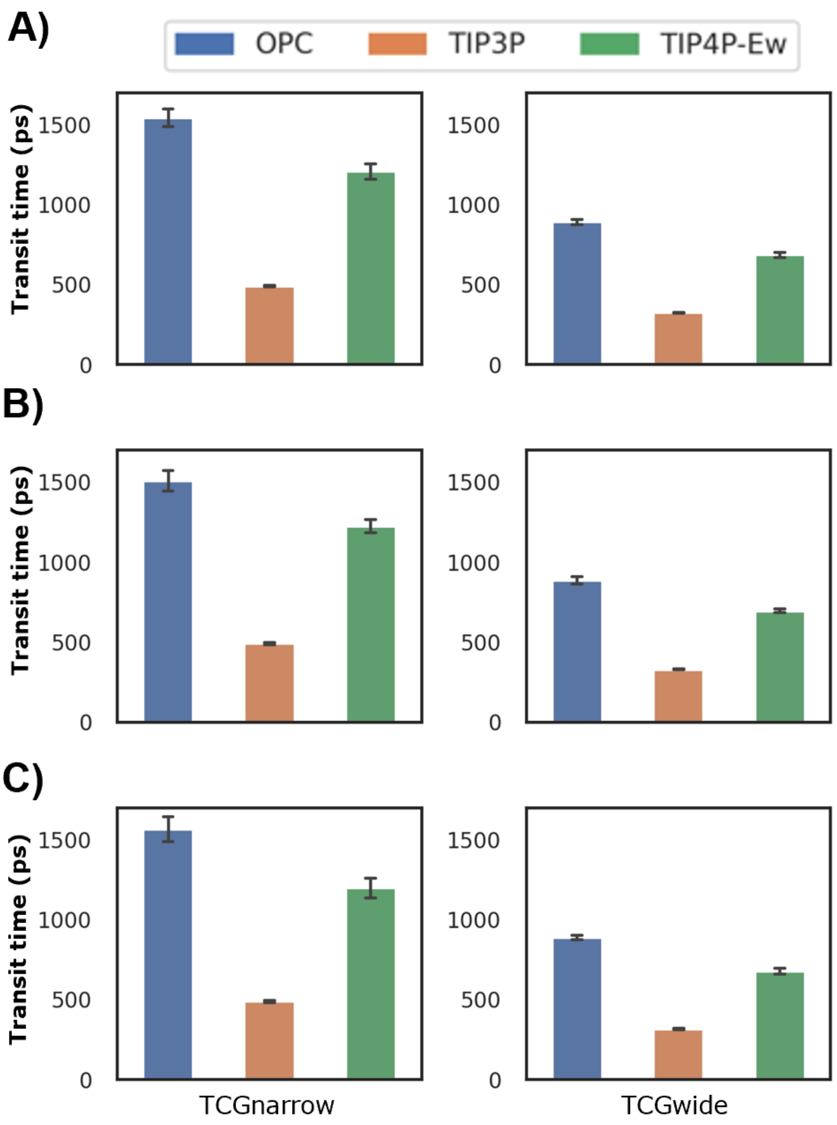

**Figure S20: Median transit time for water molecules migration across Ch2B-F tunnel for cytochrome P450.** Transit time regardless of the direction of water movement (A), water entering the active site (B), and water releasing to the bulk solvent (C). The data represent average±SEM across 5 replicated simulations.

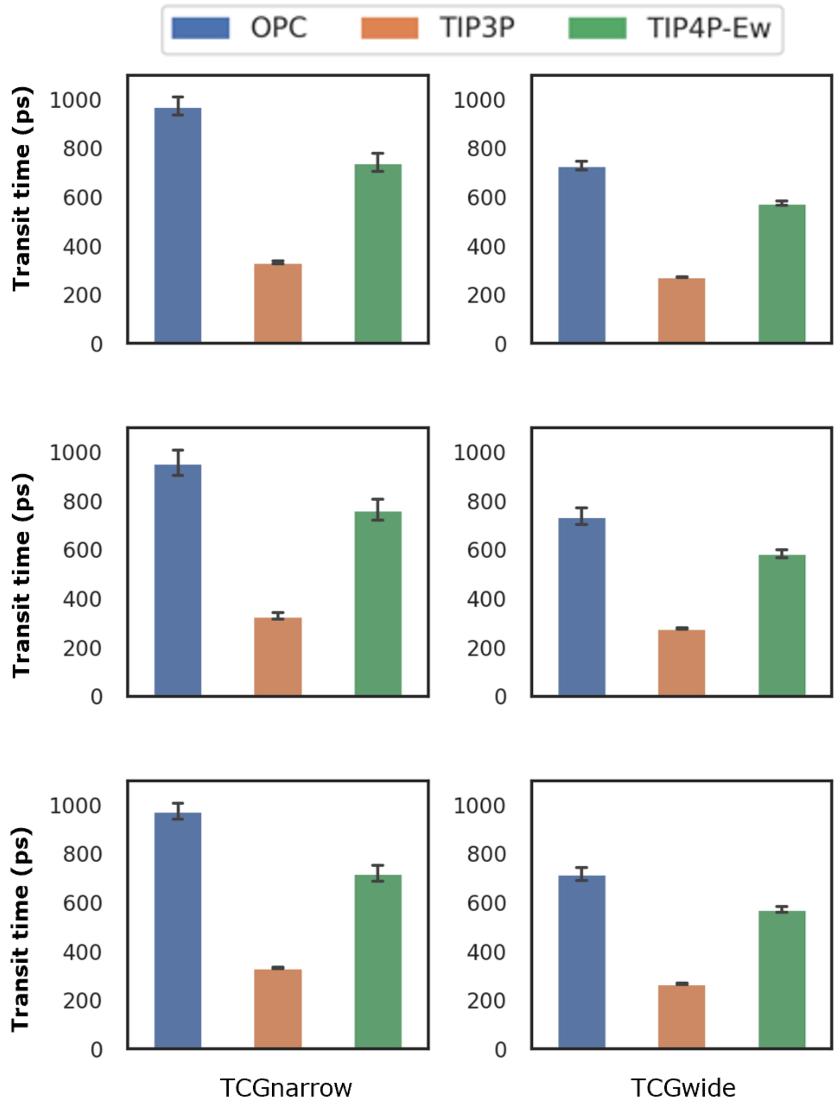

**Figure S21: Median transit time for water molecules migration across ChS tunnel for cytochrome P450.** Transit time regardless of the direction of water movement (A), water entering the active site (B), and water releasing to the bulk solvent (C). The data represent average±SEM across 5 replicated simulations.

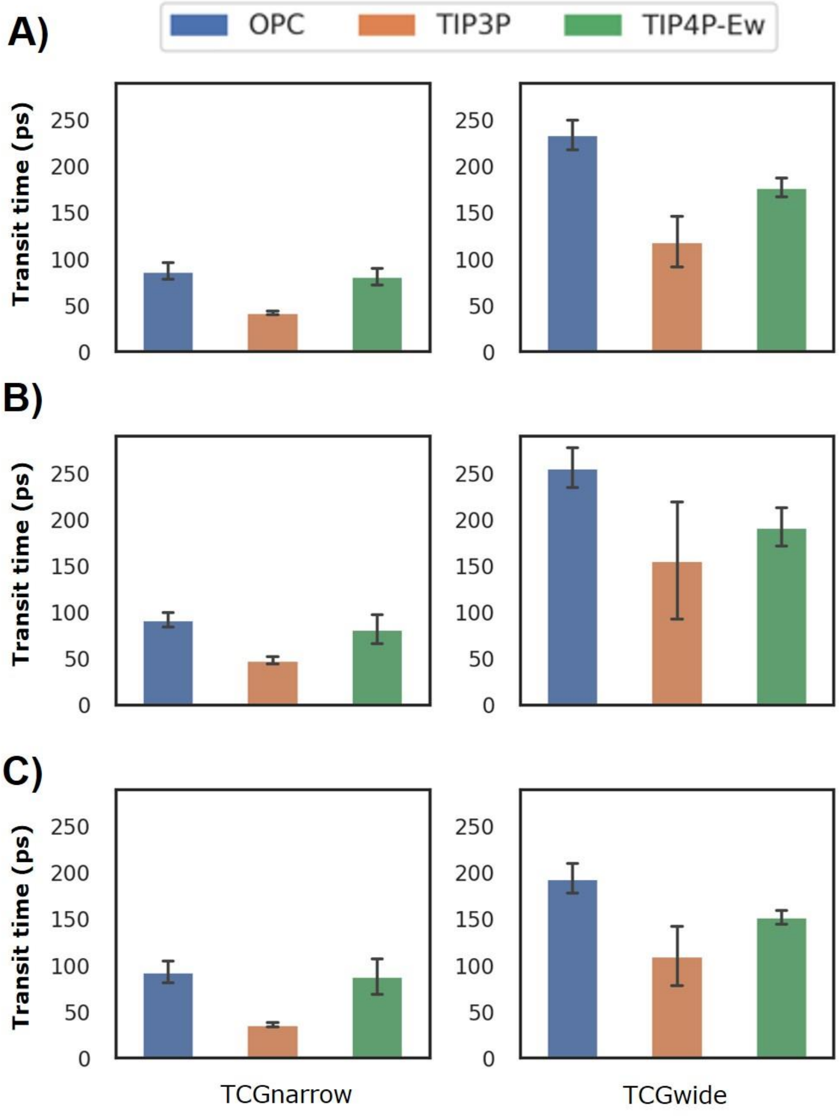

**Figure S22: Median transit time for water molecules migration across Ch2C tunnel for cytochrome P450.** Transit time regardless of the direction of water movement (A), water entering the active site (B), and water releasing to the bulk solvent (C). The data represent average±SEM across 5 replicated simulations.

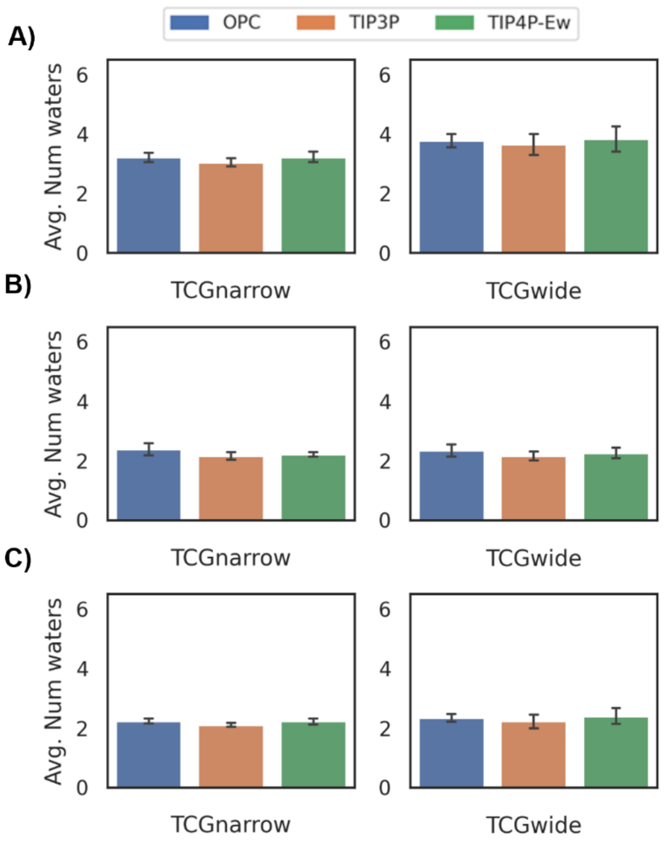

**Figure S23: Number of waters molecules concurrently transported via T1 tunnel of alditol oxidase.** The number of concurrent water molecules regardless the direction of their movement (A), entering the active site (B), and releasing to the bulk solvent (C). The data represent average±SEM across 5 replicated simulations.

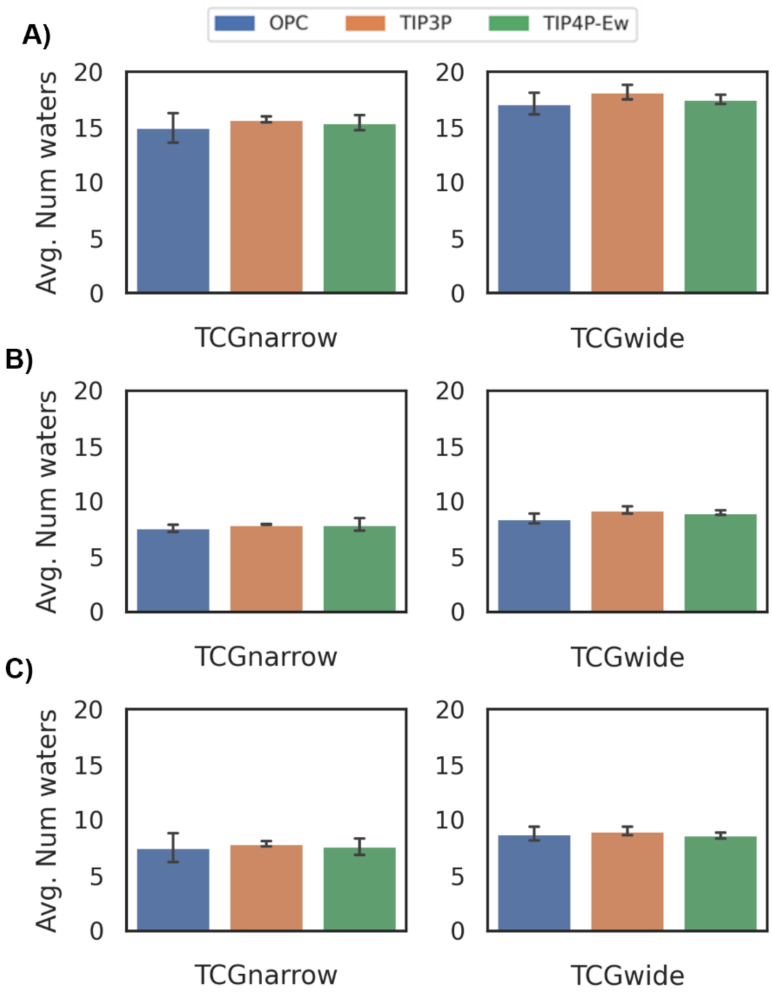

**Figure S24: Number of waters molecules concurrently transported via Ch2B-F tunnel of cytochrome P450.** The number of concurrent water molecules regardless the direction of their movement (A), entering the active site (B), and releasing to the bulk solvent (C). The data represent average±SEM across 5 replicated simulations

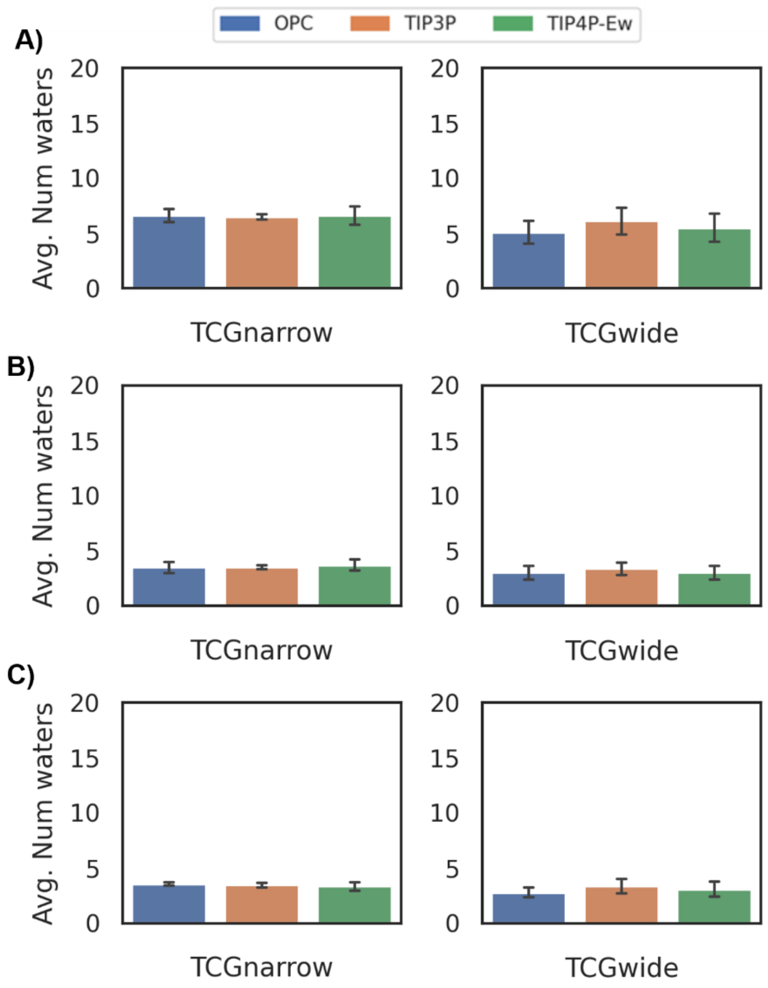

**Figure S25: Number of waters molecules concurrently transported via Ch2S tunnel of cytochrome P450.** The number of concurrent water molecules regardless the direction of their movement (A), entering the active site (B), and releasing to the bulk solvent (C). The data represent average±SEM across 5 replicated simulations

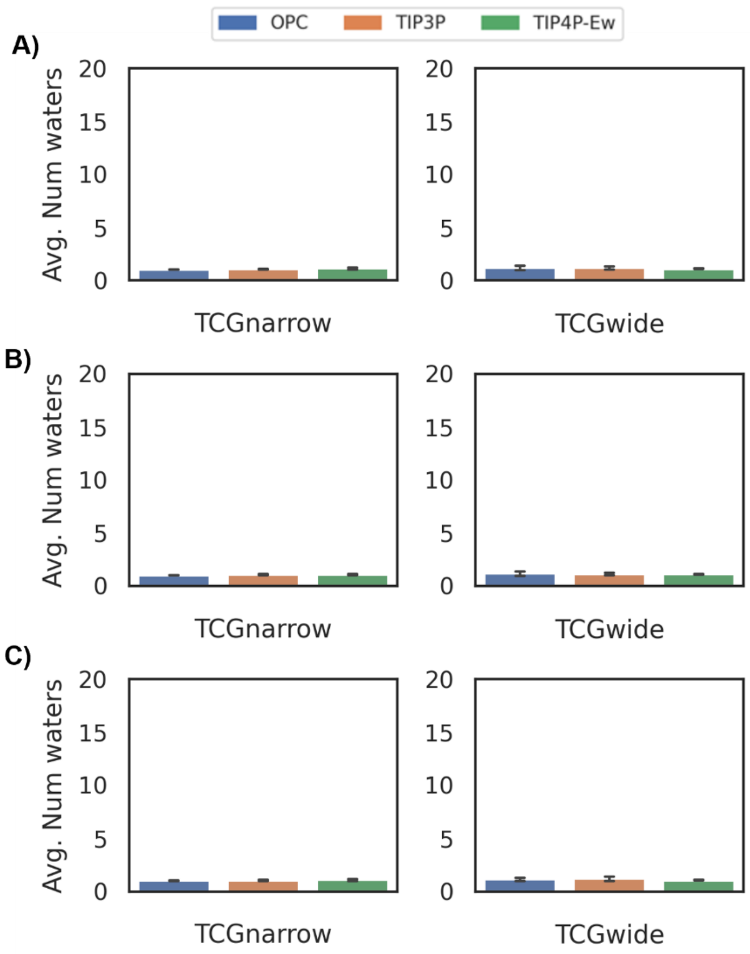

**Figure S26: Number of waters molecules concurrently transported via Ch2C tunnel of cytochrome P450.** The number of concurrent water molecules regardless the direction of their movement (A), entering the active site (B), and releasing to the bulk solvent (C). The data represent average±SEM across 5 replicated simulations.

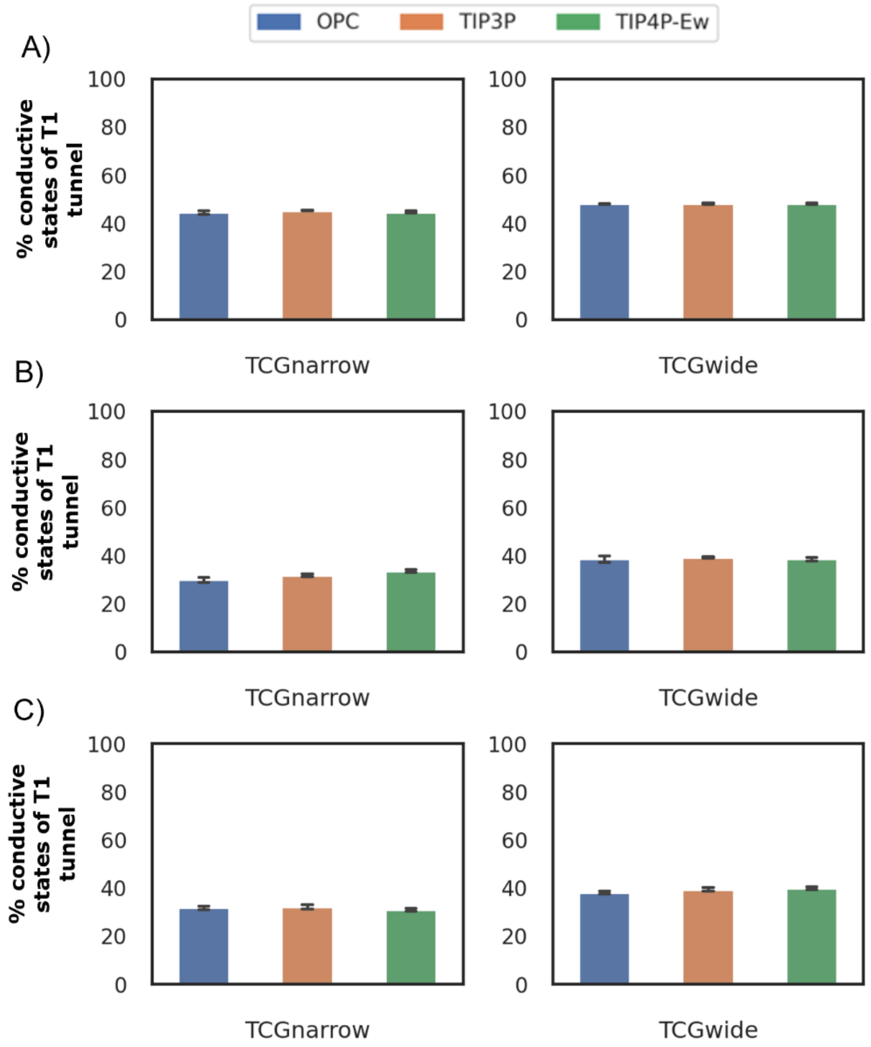

**Figure S27: Percentage of conductive states adopted by T1 tunnel in alditol oxidase for water molecules**. The percent of conductive state of waters regardless the direction of their movement (A), entering the active site (B), and releasing to the bulk solvent (C). The data represent average±SEM across 5 replicated simulations.

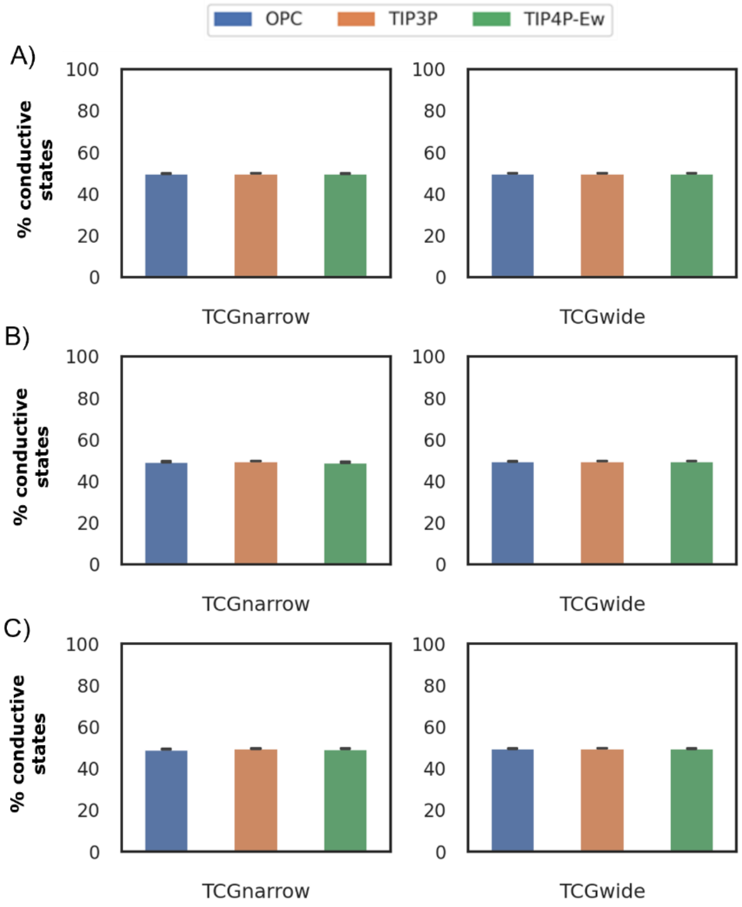

**Figure S28: Percentage of conductive states adopted by Ch2B-F tunnel in cytochrome P450 for water molecules**. The percent of conductive state of waters regardless the direction of their movement (A), entering the active site (B), and releasing to the bulk solvent (C). The data represent average±SEM across 5 replicated simulations.

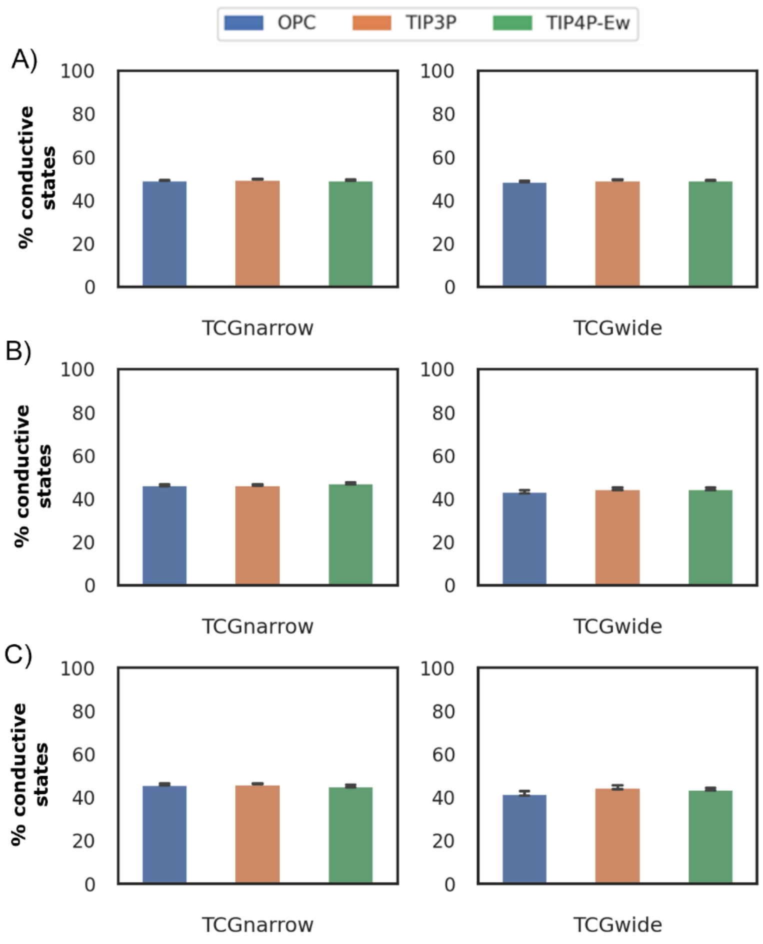

**Figure S29: Percentage of conductive states adopted by Ch2S tunnel in cytochrome P450 for water molecules**. The percent of conductive state of waters regardless the direction of their movement (A), entering the active site (B), and releasing to the bulk solvent (C). The data represent average±SEM across 5 replicated simulations.

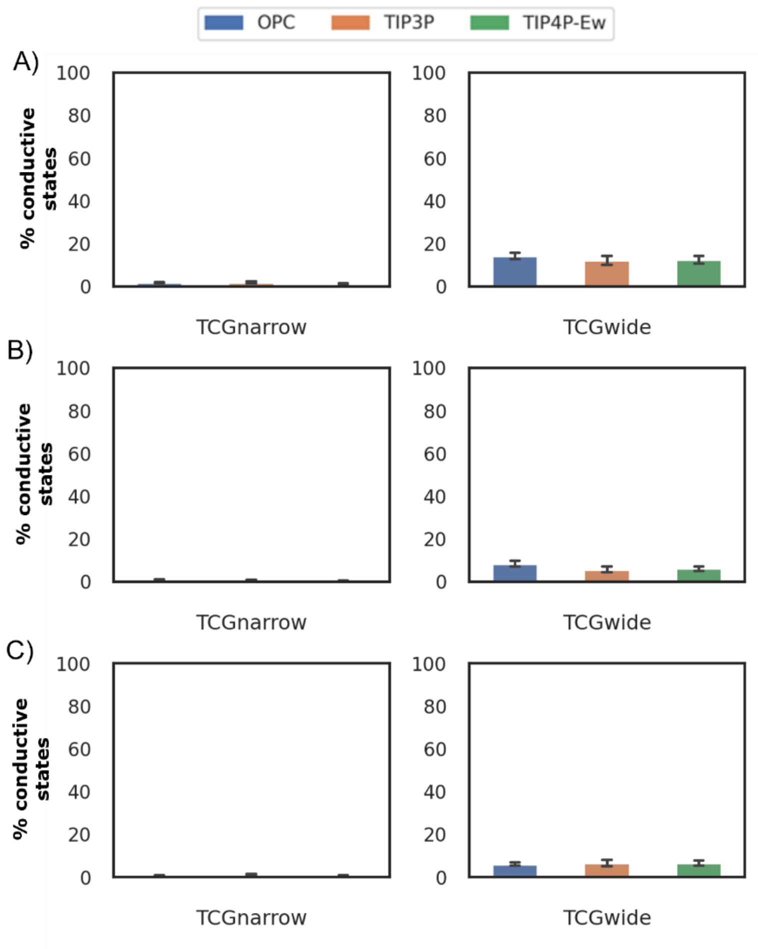

**Figure S30: Percentage of conductive states adopted by Ch2C tunnel in cytochrome P450 for water molecules**. The percent of conductive state of waters regardless the direction of their movement (A), entering the active site (B), and releasing to the bulk solvent (C). The data represent average±SEM across 5 replicated simulations.
